# Methylome inheritance and enhancer dememorization reset an epigenetic gate safeguarding embryonic programs

**DOI:** 10.1101/2021.10.10.463758

**Authors:** Xiaotong Wu, Hongmei Zhang, Bingjie Zhang, Yu Zhang, Qiuyan Wang, Weimin Shen, Xi Wu, Lijia Li, Weikun Xia, Ryohei Nakamura, Bofeng Liu, Feng Liu, Hiroyuki Takeda, Anming Meng, Wei Xie

## Abstract

Drastic epigenetic reprogramming is essential to convert terminally-differentiated gametes to totipotent embryos. However, it remains puzzling why post-fertilization global DNA reprogramming occurs only in mammals but not in non-mammalian vertebrates. In zebrafish, global methylome inheritance is however accompanied by sweeping enhancer “dememorization” as they become fully methylated. By depleting maternal *dnmt1* using oocyte microinjection *in situ*, we eliminated DNA methylation in zebrafish early embryos, which died around gastrulation with severe differentiation defects. Strikingly, methylation deficiency leads to extensive derepression of adult tissue-specific genes and CG-rich enhancers, which acquire ectopic TF binding and, unexpectedly, H3K4me3. By contrast, embryonic enhancers are generally CG-poor and evade DNA methylation repression. Hence, global DNA hypermethylation inheritance coupled with enhancer dememorization installs an epigenetic gate that safeguards embryonic programs and ensures temporally ordered gene expression. We propose that “enhancer dememorization” underlies and unifies distinct epigenetic reprogramming modes in early development between mammals and non-mammals.

## Introduction

DNA methylation plays critical roles in embryonic development, genomic imprinting, transposon silencing, and X chromosome inactivation (*1, 2*). Methylation of DNA at regulatory elements can interact with its binding proteins to exert gene repression (*3, 4*). In mammals, DNMT3A and DNMT3B can conduct *de novo* methylation (*5*). During mitosis, DNA methylation is then robustly inherited by DNMT1, facilitated by a critical co-factor UHRF1 (*6*). On the other hand, DNA demethylation is often carried out by members of the ten-eleven translocation (TET) family of enzymes (*7, 8*). Deficiency in these enzymes often leads to embryonic lethality (*9, 10*). DNA methylation at regulatory elements is considered to be associated with long-term and stable gene repression (*3*). While the numbers of promoters that are subjected to dynamic methylation are limited in the genome, DNA methylation at enhancers is highly dynamic during development and cell differentiation (*11–13*). Enhancer activation is often associated with hypomethylation, and hypermethylation at enhancer is shown to be repressive (*14, 15*). Interestingly, enhancers often remain hypomethylated even after decommission, and such state can serve as a developmental memory, as these enhancers can be activated once the corresponding TFs re-appear (*16, 17*).

In mammals, DNA methylome undergoes drastic reprogramming including global demethylation in primordial germ cells (PGCs) and pre-implantation embryos, followed by genome-wide remethylation (*18, 19*). Such reprogramming is essential in removing genomic imprints and parental memories (*20, 21*). Intriguingly, despite locus-specific reprogramming, there is no global DNA methylation reprogramming during early embryonic development in many non-mammalian vertebrates, such as zebrafish (*Danio rerio*) and *Xenopus laevis* (*22–25*). This is surprising given histone marks, by contrast, are shown to undergo global resetting in these animals (*26–29*). However, the significance of the global methylome inheritance in non-mammalian vertebrates remains elusive. In particular, previous studies showed paradoxically that zebrafish mutants deficient in zygotic *dnmt1* and *uhrf1* can survive up to a week, a stage well beyond early development as the primordial organ development is largely completed (*30–33*). While these data argue against a role of DNA methylation in embryonic development in zebrafish, it is also possible that the maternal supplies of *dnmt1* mRNAs or proteins may be sufficient to support these mutants beyond early development. In fact, knocking down uhrf1 using morpholino in zebrafish embryos (*34*) or overexpressing STELLA, a protein that can sequester UHRF1 and induce DNA demethylation, in medaka embryos (Li et al., 2018a; Mulholland et al., 2020) causes significant mortality by gastrulation. However, a similar knockdown towards *dnmt1* yielded no embryonic phenotype (*38*). It was proposed that this discrepancy may arise from methylation-independent functions of Uhrf1 or specific methylome pattern differences between *uhrf1* and *dnmt1* mutants (*38*). Therefore, whether DNA methylation is required for zebrafish embryonic development remains elusive thus far.

Despite the persisting global DNA methylation in zebrafish early embryos, marked local DNA methylation reprogramming occurs at regulatory elements during parental-to-zygotic transition. Interestingly, while promoters undergo “bidirectional” programming including both demethylation and methylation during this process (*24, 25, 27*), our previous work revealed “unidirectional” reprogramming for enhancers which become fully methylated (thus “dememorized”) either prior to fertilization for sperm, or just after fertilization for oocyte (*29*). Enhancers are not demethylated until the phylotypic stage, when Tet proteins start to be expressed (*39, 40*). However, the functional significance of such enhancer dememorization and how embryonic enhancers can operate while being unusually hypermethylated (*39, 41*) remain unanswered.

Here, we interrogated the function of DNA methylome and its inheritance in early zebrafish development, by generating *dnmt1* maternal knockdown (mKD) embryos via our recently developed technology OMIS - oocyte microinjection *in situ* (*42*). These embryos showed drastically depleted DNA methylation prior to ZGA. Importantly, these embryos failed to initiate epiboly and died around gastrulation. Careful analyses revealed defective cell differentiation, derepression of transposons, and failed establishment of Polycomb domains. Interestingly, these mutant embryos also showed widespread ectopic activation of adult enhancers and genes. Importantly, embryonic and adult enhancers show distinct CG densities and sensitivity to DNA methylation, enabling global DNA methylation as a critical epigenetic gate (“EpiGate”) to separate embryonic and adult programs. The distinct methylation sensitivity between embryonic and adult enhancers is likely sequence coded, as adult enhancers are preferentially CG-rich, while embryonic enhancers are generally CG-poor. Collectively, our study revealed that global DNA methylome inheritance is essential for early development and cell differentiation. Coupled with enhancer dememorization, DNA methylation resets an EpiGate after fertilization which safeguards embryonic programs and prevents premature firing of adult programs, to ensure temporally ordered activation of developmental programs. Furthermore, we propose that enhancer dememorization underlines epigenetic reprogramming of early development in both mammals and non-mammalian vertebrates, despite the distinct methylome reprogramming modes.

## Results

### Maternal *dnmt1* is essential for embryonic DNA methylation and development

To determine which Dnmts are responsible for maintaining embryonic DNA methylomes in zebrafish (Fig. 1A), we performed RNA-seq to oocyte, zygote, pre-ZGA embryos (4-cell and 256-cell), post-ZGA embryos (dome, a stage shortly after ZGA, and shield, a stage when gastrulation occurs) and larva (head and tail, 5 days post fertilization (dpf)). Remarkably high expression levels of *dnmt1* were found from oocyte to the 256-cell stage, indicating abundant maternal Dnmt1 (Fig. S1A). By contrast, the *de novo* Dnmts (Dnmt3/4/5/3b, the orthologs of mammalian DNMT3B, and Dnmt3aa/3ab, the orthologs of mammalian DNMT3A) show relatively low expression in zebrafish oocytes and early embryos. Our initial attempt to deplete Dnmt1 and DNA methylation by knocking down *dnmt1* in early embryos (zygotic knockdown, or zKD) or by knocking out zygotic *dnmt1* (zygotic knockout, or zKO) through crossing *dnmt1*^+/-^ heterozygotes (*31*) failed to reduce Dnmt1 protein or DNA methylation in zebrafish early embryos (Fig. 1B-C and S1B-C). Consistent with previous studies in zebrafish (*30, 31, 33*), zKD and zKO embryos can survive more than one week after fertilization (Fig. S2A). We validated zKO as the methylome analysis showed that the tail and head from zKO mutants exhibited DNA methylation loss at a later stage (larva, 5 dpf) as previously described (*31, 33*) (Fig. 1C and S1B-C). We therefore applied OMIS (oocyte microinjection *in situ*) (Fig. 1B, Methods), a method we developed recently to target maternal factors (*42*), to knockdown maternal *dnmt1* starting from oocytes rather than from zygotes (termed maternal knockdown, or mKD). Briefly, we injected *dnmt1* MO into stage III oocytes (prophase I arrested oocytes, GV stage) *in situ* while they were still kept in ovary, and the recipient female zebrafish were allowed to recover and mate with wildtype male to generate embryos naturally about 40hrs post injection. We collected embryos derived from these injected oocytes (Rhodamine B traced) at the 256-cell, dome, and shield stages, and examined the states of DNA methylation (Fig. 1B). Immunofluorescence analyses showed nearly absent signals of Dnmt1 protein or 5mC in mKD embryos from the 256-cell to shield stage (Fig. S1B, top). Low- input methylome profiling using STEM-seq (*43*) confirmed dramatic loss of global DNA methylation in mKD embryos at the 256-cell stage (from 85% to 20%), which is further exacerbated at dome (17%) and shield (7%) stages (Fig. 1C and S1C). Strikingly, *dnmt1* mKD embryos displayed severe defects and died around gastrulation (10hrs post fertilization (hpf)) (Fig. 1D-E). In wildtype embryo, cells at the margin region of blastula embryos initiate cell movements, or “epiboly”, to form dorsal- ventral axis at shield stage (*44, 45*). By contrast, *dnmt1* mKD embryos failed to initiate such epiboly, and finally died accompanied with yolk extrusion, likely caused by the ectopic stress from cell movement defects (Fig. 1D-E and S2B). *dnmt1* mKD embryos contain fewer cells especially after ZGA (Fig. S2C), indicating defects in cell proliferation and/or cell apoptosis. Therefore, maternal Dnmt1 is required for both DNA methylome inheritance and early development in zebrafish.

**Fig. 1.**
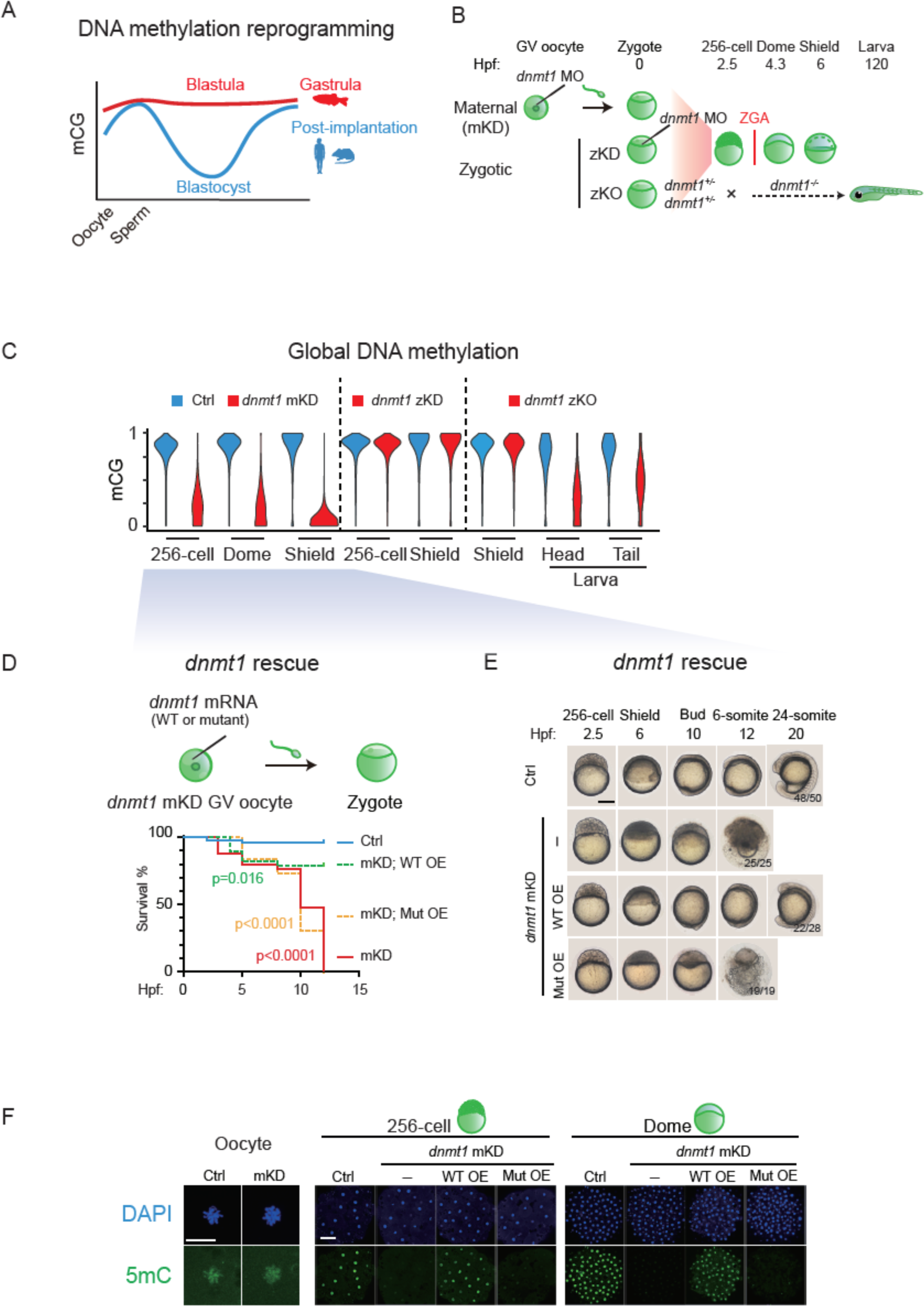
Maternal *dnmt1* is essential for embryonic DNA methylation and development. **(A)** Schematic of DNA methylation landscapes in zebrafish (red) and mammals (blue) during early development. **(B)** Schematic of *dnmt1* maternal knockdown (mKD) via oocyte microinjection *in situ*, zygotic knockdown (zKD), and zygotic knockout (zKO). Three main developmental stages were examined in this study, including the 256-cell (pre-ZGA), dome and shield stages (post-ZGA). ZGA begins around 3hrs post fertilization (hpf). zKO embryos (*dnmt1*-/-) could survive till 120hpf. Hpf, hours post fertilization. **(C)** Violin plots showing average DNA methylation levels across the genome at different developmental stages of control (blue) and *dnmt1* mKD embryos (red), zygotic knockdown (zKD) embryos (red), and zygotic knockout (zKO) (red) embryos/larvae. **(D)** Survival curve of control (blue line), *dnmt1* mKD embryos (red line), *dnmt1* mKD embryos rescued with either WT *dnmt1* mRNAs (mKD; WT OE, green dashed line) or mutant *dnmt1* mRNAs (mKD; Mut OE, orange dashed line). Log-rank test was used to calculate P-value. **(E)** Representative images of embryo phenotypes in control, *dnmt1* mKD embryos, *dnmt1* mKD embryos rescued with either WT *dnmt1* mRNAs (WT OE) or mutant *dnmt1* mRNAs (Mut OE) across different developmental stages. The numbers and ratios of embryos with a particular phenotype in each group are also shown. Scale bar, 250 μm. **(F)** Immunostaining of 5mC (green) in control and *dnmt1* mKD oocytes, 256-cell, and dome embryos, as well as *dnmt1* mKD embryos rescued by either WT (WT OE) or mutant (Mut OE) *dnmt1* mRNAs. DNA was stained with DAPI (blue). Scale bar, 50 μm.

### The loss of maternal Dnmt1 and embryonic DNA methylomes is responsible for the early lethality

We then asked if the lethality of *dnmt1* mKD embryos is caused by the loss of DNA methylation. First, co-injecting *dnmt1* morpholino with WT *dnmt1,* but not mutant *dnmt1* (both with codons modified to avoid being targeted by MO; Methods), during OMIS can efficiently rescue the epiboly delay in *dnmt1* mKD embryos (Fig. 1D-E) and restore 5mC signal at the 256-cell and dome stages, although with final levels moderately lower than the control group (Fig. 1F). Second, we examined whether mKD of *dnmt1* affects oocyte development. Both IF and low-coverage STEM-seq (due to limited numbers of mutant oocytes) analyses showed that the global DNA methylation in *dnmt1* mKD oocytes is grossly retained (Fig. 1F and S3A). In addition, the MOF (maturation, ovulation, and fertilization) (*42*) rates of oocytes were comparable between control and *dnmt1* mKD oocytes (Fig. S3B, Methods). RNA-seq of oocytes also revealed few dysregulated genes in *dnmt1* mKD oocytes compared to control oocytes (Fig. S3C). Finally, to further rule out a role of oocyte defects in the embryonic lethality caused by *dnmt1* mKD, we overexpressed mouse STELLA (also known as DPPA3), a protein that can sequester UHRF1 thus preventing the proper functions of DNMT1 to induce DNA demethylation (*35–37*), in zebrafish zygotes (instead of oocytes) (Fig. S3D-E). DNA methylation is severely impaired in *Stella* overexpressed embryos (*Stella* OE) (Fig. S3E) which showed developmental arrest at 14hrs post fertilization (somite stage) (Fig. S3F). Significant developmental delay or embryonic lethality was observed as early as shield stage and become prevalent around the somite stage (Fig. S3F). By contrast, overexpression of a mutant STELLA deficient in interaction with UHRF1 (KRR, Methods) (*35*) has little impact on 5mC level (Fig. S3E) and development (Fig. S3F). Hence, we conclude that DNA methylation is essential for early zebrafish development.

### scRNA-seq analysis revealed differentiation defects in *dnmt1* mKD embryos

We then asked how transcriptome is altered in these *dnmt1*-deficient embryos. To do so, we first performed bulk RNA-seq for control and *dnmt1* mKD embryos at the 256-cell, dome and shield stages. Globally, maternal RNA (expressed in oocytes, FPKM>10) degradation was delayed in *dnmt1* mKD embryos, but was nevertheless achieved to a large degree by shield stage (Fig. 2A). Similarly, the activation of dome-specific genes (FPKM > 10 in dome embryos and not expressed in oocytes (FPKM <5)) was delayed in mutants at dome stage, and was partially recovered at shield stage (Fig. 2A). Activation of shield-specific genes (FPKM > 10 in shield embryos and not expressed in oocytes and dome (FPKM <5)) was also partially affected (Fig. 2A). Given the cell heterogeneity of gastrula, we then performed 10× single-cell RNA-seq and profiled 19,563 and 11,852 cells from control and mKD dome embryos, respectively, and 11,288 and 20,595 cells from control and mKD shield embryos, respectively (Methods) (Fig. S4A). We confirmed the scRNA-seq results with a second replicate of lower depth (Fig. S4B). Clustering analysis for integrated data of both control and mKD embryos using Seurat (*46*) identified a total of 10 clusters, including primordial germ cells (PGCs), enveloping layer cells (EVLs), yolk syncytial layer cells (YSLs), epiblast, ectoderm, germ ring, ventral, dorsal mesoderm and dorsal margin (Fig. 2B and S4C). Most clusters were also present in *dnmt1* mKD embryos, suggesting that these embryos were able to initiate cell differentiation (Fig. S5A). However, mutant embryos showed increased epiblast cells (Fig. S5A, red arrow) and decreased ectoderm and ventral cells (Fig. S5A, blue arrow) at shield stage (Fig. S5A-B), suggesting inefficient lineage differentiation. We then analyzed cellular trajectories of control and mKD embryos in pseudotime using Monocle 2 (*47*). Cell differentiation initiates from epiblast along two major trajectories, including ectoderm (mainly located on the animal pole) and mesoderm/endoderm (mainly located in marginal zone, whose precursors are arranged along the dorsal-ventral axis) (Schier and Talbot, 2005) directions in control embryos (Fig. 2C and S5C), consistent with previous single-cell RNA-seq data in zebrafish early embryos (*48, 49*). By contrast, *dnmt1* mKD shield embryos exhibited multiple branches from epiblast that led to a mixture of differentiated cells and undifferentiated epiblast, suggesting aberrant differentiation programs. Consistently, downregulated genes of each cell cluster are overwhelmingly enriched for developmental genes and genes related to gastrulation and cell movement (Fig. 2D and S5D-E). Several lineage markers showed incorrect spatial expression patterns, including *tph1b*, *frzb* (for dorsal margin) and *eve1*, *dld* (for ventral margin) (Fig. S6). Notably, developmental genes, including some that are typically expressed in adult tissues such as *ntsr1*, *ncaldb*, *atxn1b*, and *lrrtm1*, were also enriched in upregulated genes in mKD embryos (Fig. 2D and S6) (discussed in detail later). Hence, scRNA-seq analysis revealed widespread differentiation defects in *dnmt1* mKD embryos.

**Fig. 2.**
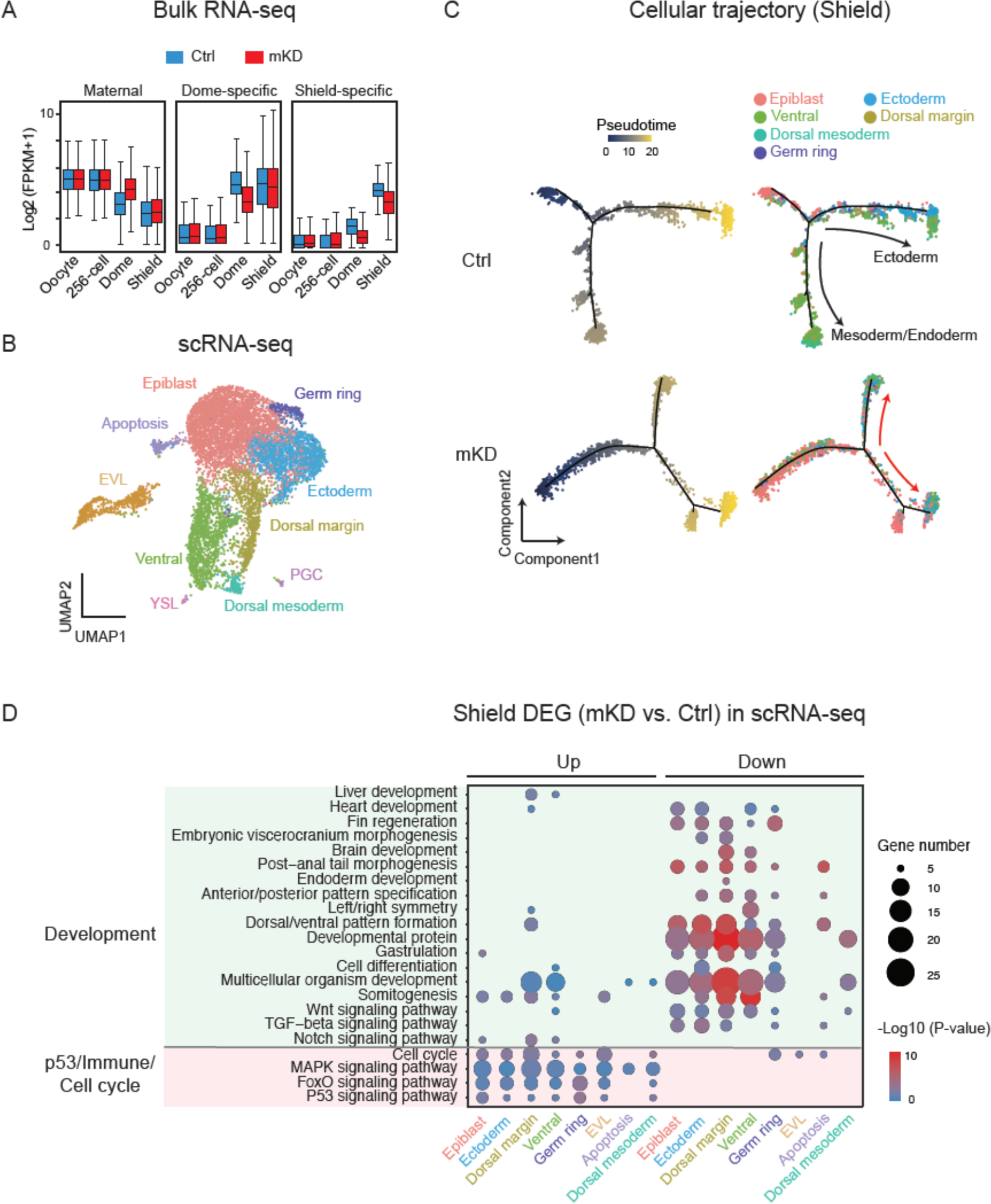
Bulk and single-cell RNA-seq revealed developmental defects in mKD embryos. **(A)** Distribution of RNA levels of maternal genes (left), dome-specific genes (middle) and shield-specific genes (right) across different developmental stages of control (blue) and *dnmt1* mKD (red) embryos. **(B)** Projection of cells with UMAP for control and *dnmt1* mKD embryos at dome and shield stages. Cells are colored by clusters. **(C)** Pseudotime trajectory of control and *dnmt1* mKD embryos at shield stage. Cells were ordered from epiblast to ectoderm or mesoderm and endoderm, and colored by pseudotime (left) or clusters in **b** (right). Red arrows indicate abnormal cell differentiation branches. **(D)** Bubble plot showing enriched GO terms for differentially expressed genes (DEGs) between control and *dnmt1* mKD embryos of each cluster at shield stage. Top enriched terms include development and *p53* dependent apoptosis, immune response and cell cycle related genes. Size of circle encodes gene number; color of the circle indicates -log10(P-value).

### *dnmt1* mKD embryo lethality is partially contributed by transposon-derepression triggered immune response and *p53*-mediated cell apoptosis

Notably, upregulated genes in *dnmt1* mKD embryos are also enriched for *p53* signaling, cell cycle and immune response (Fig. 2D and S5E), consistent with the observation in hypomethylated *dnmt1* zKO zebrafish larva (> 3 days post fertilization) where innate immune response and *p53*-mediated cell apoptosis are often activated upon the loss of DNA methylation due to the activation of transposons (*50*). Indeed, using WT and *p53* null female zebrafish (*51*) (Fig. S7A-B), we observed increased cell apoptosis in *dnmt1* mKD *p53*^+/+^ embryos but not in *dnmt1* mKD *p53*^-/-^ embryos (Fig. S7C). The *dnmt1* mKD *p53*^-/-^ embryos could now initiate epiboly movements and form dorsal-ventral axis, although with a delayed kinetics. However, these embryos eventually died around 12hpf (Fig. S7B), suggesting that *p53*-dependent cell apoptosis is only partially responsible for embryo lethality in *dnmt1* mKD embryos. Furthermore, analysis of the total RNA-seq data revealed derepression of several classes of TEs, in particular, LTRs and their subfamily *gypsy*, in mKD embryos at dome stage, which is further exacerbated at shield stage (Fig. S7D-F). This is accompanied by increased genome instability, as manifested by increased phosphorylated H2A.X signals, a marker of DNA double strand breaks (DSBs) (*52*) (Fig. S8A). Similar observation was made for STELLA overexpressed zebrafish embryos (Fig. S8B). We then asked whether the induced DSBs may be partially responsible for the lethality of DNA methylation-deficient embryos, by treating embryos with Foscarnet (FOS, Methods), an inhibitor of reverse transcriptases and polymerases and an effective antiviral agent (*53*). We used *Stella* OE embryos in this experiment as *dnmt1* mKD using OMIS could not produce large numbers of embryos required for drug treatment and statistical analysis. Although effectively eliminated DSB from the *Stella* OE embryos, FOS treatment however could not rescue their developmental defects (Fig. S8B-C). These data indicate that the transcription of transposon itself, but not its retrotranscribed DNA or the subsequent DNA damage, may trigger developmental defects in *dnmt1* mKD embryo. Phosphorylation of Tbk1 (pTbk1) acts as an indicator of viral sensor signaling activation (*54*). The treatment of BX795, an inhibitor of pTbk1 that can repress interferon-response genes in zebrafish (*50*), could rescue a small fraction (9/111, 8.1%) of *Stella* OE embryos (Fig. S8C). Therefore, we conclude that *p53*-dependent cell apoptosis and pTbk1-mediated immune response, likely triggered by transposon derepression, partially contribute to developmental defects in *dnmt1* mKD embryos.

### The loss of promoter DNA methylation is responsible for part, but not all, of gene derepression in *dnmt1* mKD embryos

Next, we investigated how the transcription defects in mKD embryos are related to the loss of DNA methylation. As the lingering maternal transcripts may mask embryonic transcription, we excluded “maternal genes” (FPKM>5 in oocytes) from the analysis. Given the repressive roles of DNA methylation, we focused on upregulated genes and interrogated their expression and promoter methylation states (Fig. 3A-B). We first examined shield stage embryos, where gene upregulation is more evident in mutants (Fig. 3A-B). About 399 upregulated genes show the loss of DNA methylation at promoters, which mainly function in plasma membrane and fibronectin (involved in cell movements likely related to epiboly), cell migration and G-protein coupled receptor (Fig. 3B, red). Interestingly, the rest 260 upregulated genes showed no significant changes of DNA methylation (Fig. 3B, orange). In fact, 98% of these promoters are hypomethylated in both control and mKD embryos. These genes are preferentially enriched for fin morphogenesis, blood vessel development and CNS (central nervous system) development (Fig. 3B, orange), which appear to be more related to adult tissue development. Examples of adult development related genes, including *ntsr1*, *atxn1b*, *ncaldb*, *lrrtm1*, were also observed in scRNA-seq data (Fig. S6). For convenience, we termed these two groups as “promoter methylation dependent” and “promoter methylation independent” genes. A similar analysis of dome stage embryos revealed 130 promoter methylation dependent and 54 promoter methylation independent genes, with 55.4% of promoter methylation dependent genes overlapping those at shield stage (Fig. S9A-B). In sum, promoter DNA methylation loss is likely responsible for some, but not all, derepressed genes.

**Fig. 3.**
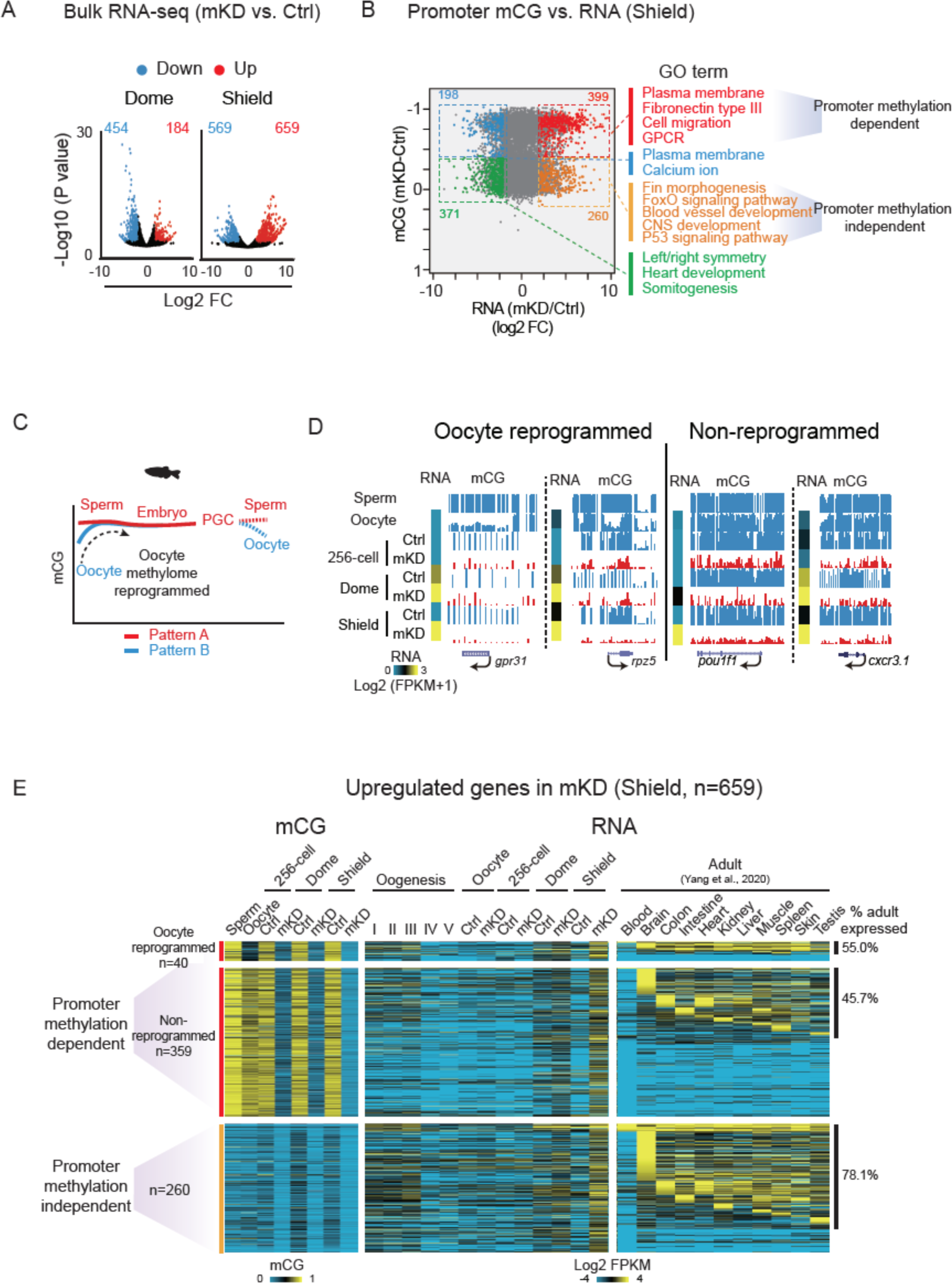
Promoter DNA methylation dependent and independent gene derepression in mKD embryos. **(A)** Volcano plots showing gene expression (log2(FPKM + 1)) changes between control and *dnmt1* mKD embryos at dome and shield stages. Red dots indicate upregulated genes; blue dots indicate downregulated genes. The numbers in corresponding colors indicate counts of dysregulated genes. **(B)** Scatter plots comparing alteration of gene expression (log2FC (mKD/Ctrl)) and promoter mCG between control and *dnmt1* mKD embryos at shield stage. Red and orange dots indicate promoter DNA methylation dependent and independent upregulated genes, respectively; blue and green dots indicate downregulated genes with decreased and constant promoter DNA methylation, respectively. The numbers of dysregulated genes and enriched GO terms in corresponding group (color coded) are also shown. **(C)** Schematic of DNA methylation reprogramming from gametes to the next generation in zebrafish. Sperm, early embryo and PGC exhibit highly similar methylomes (pattern A) (Jiang et al., 2013; Potok et al., 2013; Skvortsova et al., 2019). Oocyte has a distinct methylome (pattern B) that will be reprogrammed to pattern A after fertilization. **(D)** UCSC genome browser snapshots showing promoter mCG in sperm, oocyte, 256-cell, dome and shield stage embryos. RNA of the 256-cell, dome and shield stage embryos are also shown (heat map). **(E)** Heat maps showing promoter mCG and RNA expression of promoter DNA methylation dependent or independent genes in **b** (n=659) across developmental stages of oogenesis, early embryos and adult tissues (*56*). Promoter DNA methylation dependent genes were further classified into oocyte reprogrammed (mCG (dome or shield - oocyte) > 0.4) and non-reprogrammed (mCG (dome or shield - oocyte) <= 0.4) groups based on the mCG levels in oocyte and early embryos. The ratios of tissue expressed genes (FPKM >5) in each group are also shown, and statistical significance for the enrichment was assessed with one-sided Fisher’s exact test.

Notably, despite the inheritance of global DNA methylome after fertilization, promoter-specific DNA methylation reprogramming does occur in zebrafish early embryos (*24, 25*). In particular, promoter methylation on the maternal genome is reconfigured (including both methylation and demethylation) to a pattern that is similar to that of the paternal genome (Fig. 3C) (*24, 25*). Such “sperm-like” methylome also persists to PGCs (*55*). The significance of such reprogramming, however, remains elusive. We identified 842 promoters that are hypomethylated in oocytes but become methylated after fertilization (Fig. S9C). About 41.0% of them are expressed during oogenesis or PGCs, and 32.5% are expressed during embryonic development or in adult tissues (Fig. S9C-D). However, only 4.8% (n=40 out of 842) of these “reprogrammed” genes are derepressed in mKD embryos by shield stage. Therefore, it appears that the function of such oocyte-to-embryo reprogramming is not just restricted to gene repression during the imminent ZGA. Rather, these data raise an interesting possibility that such reprogramming may restore promoter methylation to a “ground” state to facilitate both ZGA and future development.

We then asked when these derepressed genes normally express in WT zebrafish. Using a collected RNA-seq data from a total of 11 adult tissues (*56*), we found that 46.6% of promoter methylation dependent group are expressed in at least one adult tissue (55.0% for oocyte reprogrammed and 45.7% for non-reprogrammed, Fig. 3E). The ratio of tissue expressed genes is however substantially higher for promoter methylation independent group (78.1%, Fig. 3E, also compared to 53.3% of random genes). This also echoes GO term analysis which showed that promoter methylation independent group genes are preferentially involved in later development (such as “fin morphogenesis”, “blood vessel development” and “CNS development”) (Fig. 3B). Overall, 389 (59.0%) upregulated genes are expressed in adult tissues. To confirm that these genes are preferentially expressed later during development, we examined their expression using scRNA-seq data from 4hrs to 24hrs post fertilization of WT embryos (*49*). We found only 19.3% (75 of 389) are expressed prior to 24hpf (phylotypic stage), while the rest 80.7% (311) are activated at least after 24hpf (Fig. S9E). Therefore, these data indicate adult programs are aberrantly activated in *dnmt1* mKD embryos.

### DNA methylation is required for the proper establishment of Polycomb domains

We then investigated what may contribute to promoter methylation independent gene derepression in *dnmt1* mKD embryos. Histone modifications undergo extensive global loss and re-establishment during early zebrafish development (*26–29*). In particular, Polycomb domains, marked by the repressive marks H3K27me3 and H2AK119ub, are established around ZGA (*57–59*). These marks, deposited by PRC2 and PRC1, respectively, are critical repressors for key developmental genes (*60*). Thus, we collected *dnmt1* mKD embryos at dome and shield stages and performed H3K27me3 and H2AK119ub CUT&RUN (*61*). Strikingly, H3K27me3 and H2AK119ub re-establishment are severely impaired in *dnmt1* mKD embryos (Fig. 4A-B and S10A). About 51.7% (420/808) promoter H3K27me3 peaks are lost or strongly reduced in *dnmt1* mKD embryos. Interesting, the rest (48.3%, 388/808) appear to be largely intact (Fig. 4B). These two groups show similar enrichment for developmental genes (Fig. S10B), although their promoters appear to enrich for distinct TF motifs (Fig. 4B). Although both classes of promoters are enriched for CGs, the “retained” group show higher CG levels at promoters (Fig. 4B-C), consistent with the notion that CG-rich sequences can recruit Polycomb (*62*). The fact that these marks at some, but not other, genes are affected suggest that this is not simply due to developmental delay. IF analysis also excluded a possibility of global H3K27me3 decrease in mKD embryos (Fig. S10C). A similar trend was observed for H2AK119ub (Fig. 4A-B). This result is reminiscent of the observation in mouse ESCs, where the loss of global DNA methylation leads to decrease of promoter H3K27me3 (*63, 64*), and supports the model that the absence of DNA methylation in the genome elsewhere may allow the spreading and/or recruitment of Polycomb, which in turn dilutes Polycomb and H3K27me3 away from promoters (Bartke et al., 2010; Brinkman et al., 2012; Hagarman et al., 2013; Reddington et al., 2013; Wu et al., 2010). Nevertheless, the decreased H3K27me3 and H2AK119ub did not cause apparent widespread gene derepression (Fig. 4C-D), as only 38 (9.1%) genes that lost promoter H3K27me3 showed moderate derepression, including *bdnf*, *pax7b* and *tat* (Fig. 4D). Therefore, these data demonstrate that DNA methylation is crucial for proper establishment of Polycomb domains at developmental gene promoters in early zebrafish embryos. However, the loss of repressive marks, such as H3K27me3 and H2AK119ub, cannot fully explain promoter methylation-independent gene derepression in *dnmt1* mKD embryos.

**Fig. 4.**
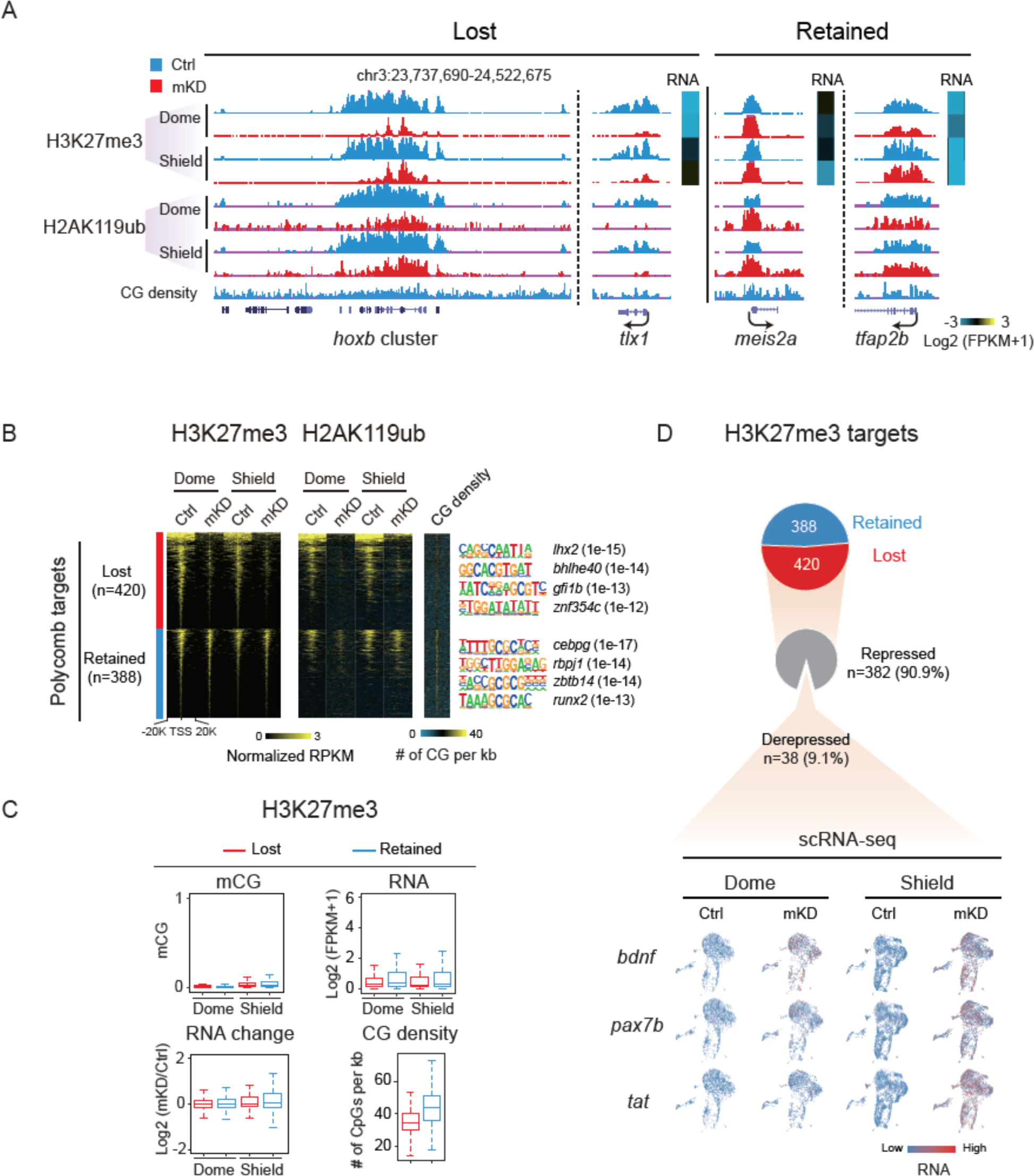
*dnmt1* mKD results in defective Polycomb domain establishment. **(A)** A UCSC genome browser snapshot showing H3K27me3 and H2AK119ub of control (blue) and *dnmt1* mKD (red) embryos at dome and shield stages. Heat map showing RNA expression of related genes. **(B)** Heat maps showing the normalized H3K27me3 and H2AK119ub enrichment around promoters (TSS±20kb) of Polycomb target in control (blue) and *dnmt1* mKD (red) embryos at dome and shield stages. CG density and enriched TF motifs are shown too (right). **(C)** Box plots showing promoter mCG, RNA and CG density levels for two clusters of Polycomb targets (lost (red) and retained (blue), defined in **b**) at dome and shield stages of control (top). RNA changes between control and *dnmt1* mKD embryos (RNA change) are also shown (bottom left). **(D)** Pie chart showing the percentages of Polycomb target genes with lost or retained H3K27me3 in *dnmt1* mKD embryos at shield stage, and among “lost” group genes the percentages of genes that are derepressed (log2FC (mKD/Ctrl) > 2) and remain repressed (grey) (top). The feature plots of scRNA-seq show RNA expression of example derepressed genes in control and *dnmt1* mKD embryos at dome and shield stages (bottom). Each dot indicates one cell. Red, high expression; blue, low expression.

### Loss of DNA methylation results in aberrant activation of adult enhancers with ectopic H3K4me3 and TF binding

Besides promoters, distal regulatory elements, such as enhancers, play crucial roles in gene regulation (*69*). To ask whether they may play a role in gene derepression in *dnmt1* mKD mutants, we profiled histone marks H3K4me3, H3K4me1, H3K27ac (Methods), and chromatin accessibility using ATAC- seq (*70, 71*). Strikingly, we found that H3K4me3, a typical permissive promoter mark, is highly dynamic upon the loss of DNA methylation especially in distal regions (Fig. 5A-B). While the overall numbers of promoter H3K4me3 peaks showed a moderate decrease from 19,385 to 16,913, the distal H3K4me3 peaks increase dramatically from 2,099 to 13,457 (Fig. 5A). As a result, the ratio of distal H3K4me3 peaks among all H3K4me3 peaks increases substantially from 9.8% in control to 44.3% in mKD embryos. Even after excluding weak distal H3K4me3 peaks (normalized RPKM < 0.5 in both control and mKD samples), there are still 7,580 distal H3K4me3 peaks left. To rule out the possibility that these regions may be unannotated promoters, we further excluded distal H3K4me3 peaks that overlap with any H3K4me3 peaks (promoter mark) in adult tissues (*56*) and unmethylated regions in the 256-cell embryos (when all enhancers are presumably methylated, leaving only promoters unmethylated) (*29*) (Methods). This still yielded a total of 6,380 distal H3K4me3 peaks. Such widespread distal H3K4me3 is unique to mutant embryos, as we only identified 32 distal H3K4me3 peaks specifically in control embryos, and 298 peaks present in both control and mKD embryos (Fig. 5C). To understand the nature of these ectopic distal H3K4me3 sites, we mapped the states of DNA methylation, H3K27ac, H3K4me1, open chromatin (ATAC-seq) and CG density in different groups based on whether a distal H3K4me3 peak is lost, ectopically acquired or retained in mutant (Fig. 5C). Indeed, ectopic distal H3K4me3 sites also showed substantially increased H3K27ac, H3K4me1, and chromatin accessibility, as manifested globally (Fig. 5C) and also at individual genes (Fig. 5D), indicating enhanced regulatory activities. Notably, a fraction of these regions (11.4%) also showed H3K27ac in control embryos (dome and shield), suggesting that these elements are likely active in WT embryos and their activities further increased in mutants (discussed later).

**Fig. 5.**
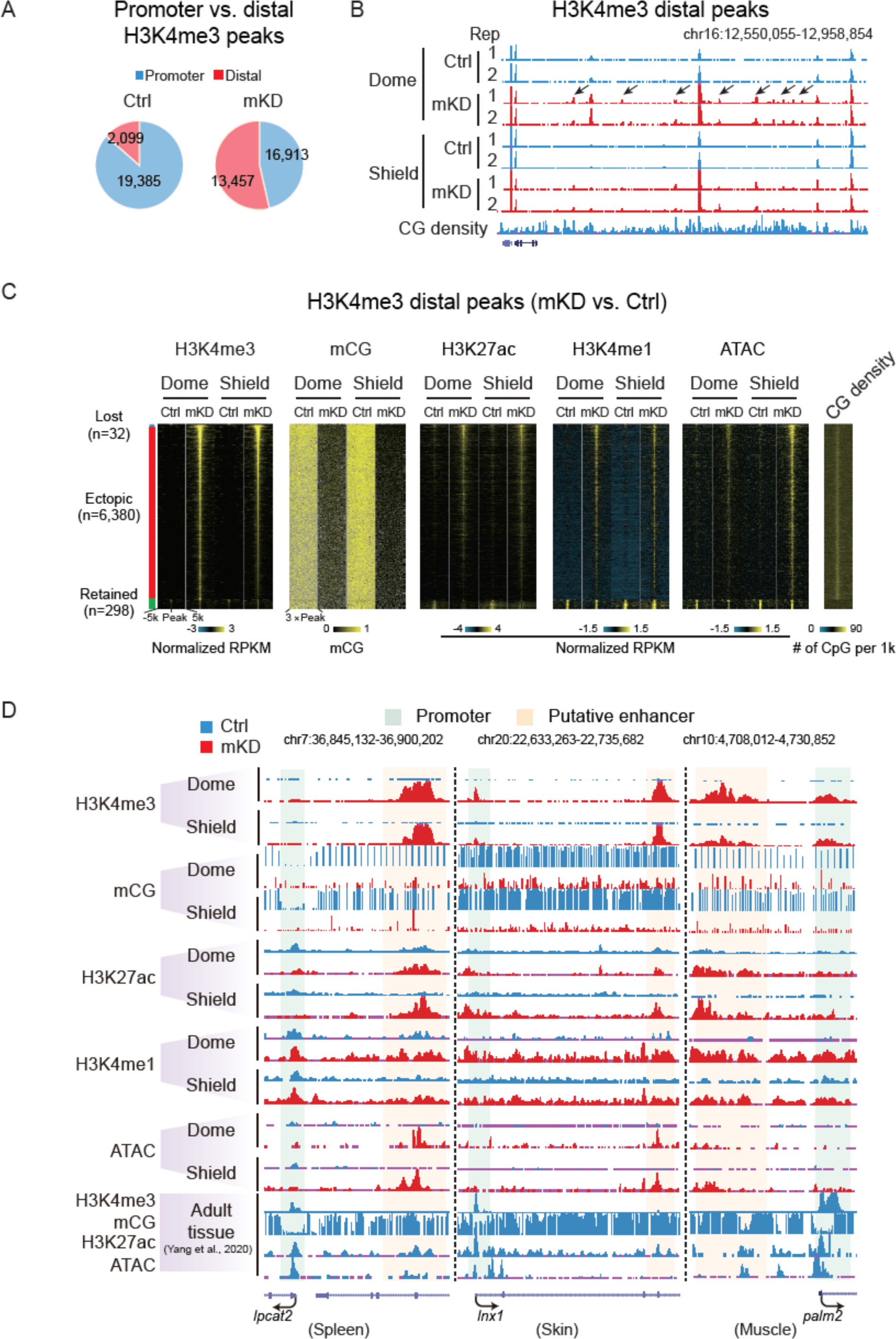
Loss of DNA methylation causes ectopic activation of putative adult enhancers and aberrant acquisition of distal H3K4me3 in early embryos. **(A)** Pie charts showing distributions of promoter (blue) and distal (red) H3K4me3 sites in control and mKD embryos at shield stage. **(B)** UCSC genome browser snapshot showing H3K4me3 at dome and shield stages of control (blue) and *dnmt1* mKD (red) embryos, and CG density. Arrows indicate ectopic H3K4me3 sites. **(C)** Heat maps showing distal H3K4me3, mCG, H3K27ac, H3K4me1, open chromatin (ATAC-seq) and CG density in either control or *dnmt1* mKD embryos. Peaks were classified into three clusters based on the dynamics of distal H3K4me3: lost (H3K4me3 lost in *dnmt1* mKD embryos, blue), ectopic (H3K4me3 acquired in *dnmt1* mKD embryos, red), and retained (H3K4me3 present in both control and mKD, green). **(D)** UCSC genome browser snapshots showing H3K4me3, mCG, H3K27ac, H3K4me1, and open chromatin (ATAC-seq) at dome and shield stages of control (blue) and *dnmt1* mKD (red) embryos, and adult tissues (Spleen, left; Skin, middle; Muscle, right) (*56*). Green shadow indicates promoter, and orange shadow indicates putative enhancer.

Given H3K27ac, accessible chromatin, and, in particular, H3K4me1, are considered as hallmarks for enhancers (*69*), we then asked if these elements are possibly enhancers. As the majority of them are not active in control early embryos, we examined their chromatin states in 11 adult tissues (*56*).

Encouragingly, 24.2% (1,547/6,380) are marked by H3K27ac, which marks active enhancers (*72*), in at least one adult tissue (Fig. 6A). Enhancers can also stay in poised or decommissioned states, which no longer bear H3K27ac, but are still marked by accessible chromatin or DNA hypomethylation (*16, 17, 73*). Strikingly, more than half (54.4%, 3,472/6,380) of these ectopic distal H3K4me3 overlap with lowly methylated regions (LMRs) identified in tissues (Fig. 6B). The ectopic H3K4me3 peaks also precisely align with the center of LMRs. Overall, we found that among all 6,380 ectopic distal H3K4me3 sites, the majority (63.1%, 4,026/6,380) overlap with putative enhancers that are either active (marked by H3K27ac) or poised/decommissioned (marked by LMRs or accessible chromatin) in at least one of 11 adult tissue lineages (Fig. 6C). By contrast, only a small fraction (11.4%) overlaps with early embryonic putative enhancers (dome and shield stages). The rest 1,628 (25.5%) peaks did not overlap with adult enhancers or embryonic enhancers (Fig. 6C, “Rest”). We expect that this number would further decrease when adding additional tissue types and developmental stages. Finally, we asked if the presumably derepressed enhancers are within the proximity of derepressed genes. Indeed, these putative enhancers are closer to upregulated genes but not to downregulated genes or non-DEGs (differentially expressed genes) (Fig. 6D). Hence, these data suggest that these ectopic distal H3K4me3 sites preferentially occupy putative adult enhancers, which are linked to derepression of adult genes.

**Fig. 6.**
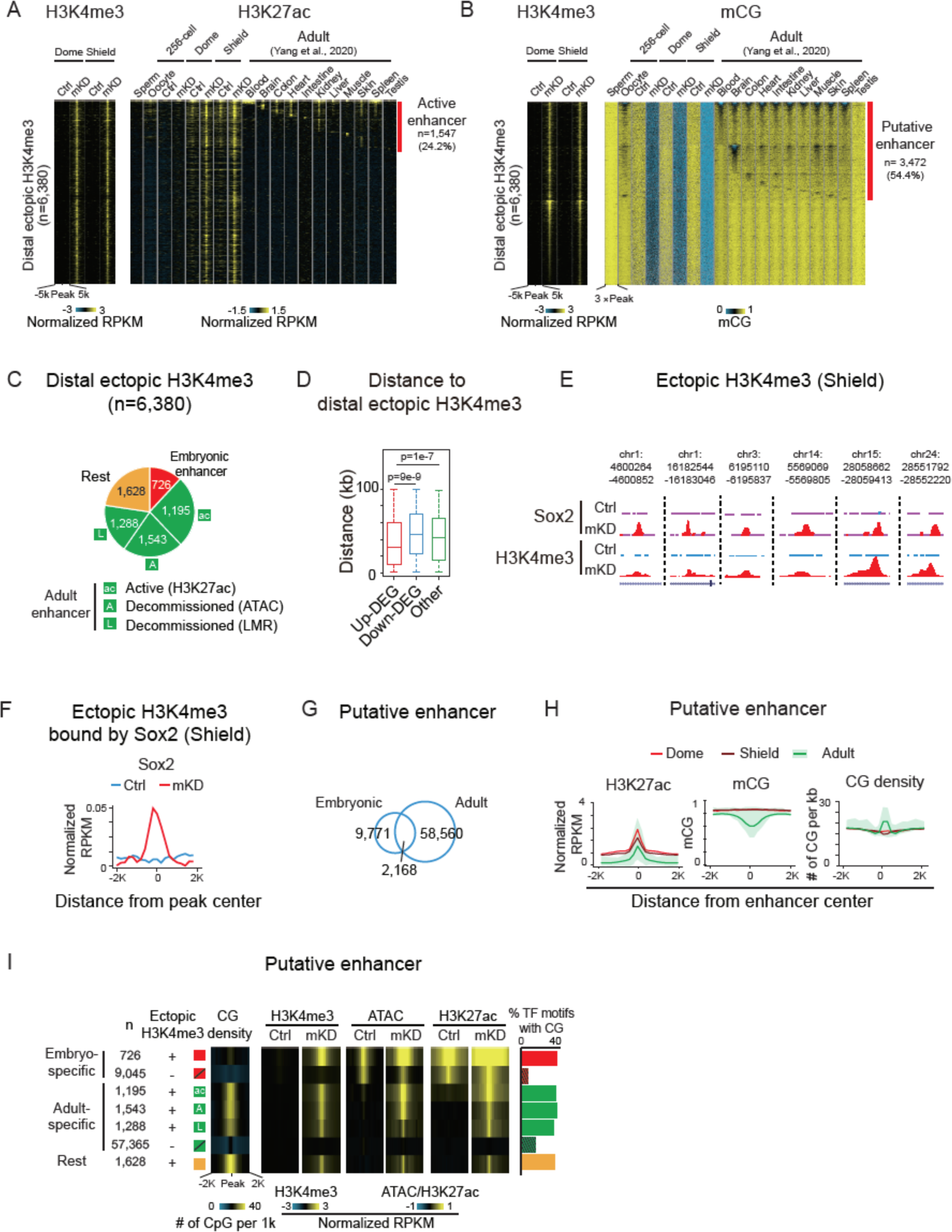
DNA methylation resets embryonic and adult programs in zebrafish through blocking high CG density adult enhancers. **(A)** Heat maps showing distal H3K4me3 and H3K27ac around distal ectopic H3K4me3 peaks in either control or *dnmt1* mKD embryos, and WT sperm, oocyte, and adult tissues (*56*). Peaks with H3K27ac signals in adult tissues are defined as active enhancers (red). **(B)** Heat maps showing distal H3K4me3 and mCG around distal ectopic H3K4me3 peaks in either control or *dnmt1* mKD embryos, and WT sperm, oocyte, and adult tissues (*56*). LMRs (lowly methylation regions) in adult tissues are defined as putative enhancers (red). **(C)** Pie chart showing distribution of distal ectopic H3K4me3 overlapping with embryonic enhancers (red), active adult enhancers (green, “ac”, defined by distal H3K27ac), decommissioned adult enhancers (green, “A”, defined by ATAC-seq without H3K27ac; “L”, defined by LMRs without H3K27ac), and the rest ectopic H3K4me3 sites (“rest”, orange). **(D)** Box plot showing distance from center of distal ectopic H3K4me3 peaks to upregulated (red), downregulated (blue), and non-DEGs (green) in *dnmt1* mKD embryos at shield stage. **(E)** UCSC genome browser snapshots showing Sox2 binding and H3K4me3 at shield stage in control (blue) and *dnmt1* mKD (red) embryos. **(F)** Line chart showing average Sox2 binding distribution around ectopic H3K4me3 regions acquiring Sox2 at shield stage in control (blue) and *dnmt1* mKD (red) embryos. **(G)** Venn diagram showing overlap between embryonic (dome and shield combined) and adult enhancers (11 adult tissues combined) (*56*). **(H)** Line charts showing H3K27ac, mCG, and CG density in control embryos at dome (red) and shield (dark red) stages, and adult tissues (*56*) (green line, average value of all tissues; green shade, range of all tissues) around putative enhancer regions defined at each stage. **(I)** Heat maps showing average CG density, H3K4me3, open chromatin (ATAC-seq) and H3K27ac (*56*) in various classes of putative enhancers in either control and mKD embryos. Definitions of putative enhancer classes are shown on the left: red box, embryonic-specific enhancer with distal ectopic H3K4me3; crossed red box, embryonic-specific enhancer without distal ectopic H3K4me3; green boxes with “ac”, “A”, and “L”, three clusters of adult enhancers with distal ectopic H3K4me3 defined in **c**; crossed green box, adult enhancers without distal ectopic H3K4me3; orange box, the rest distal ectopic H3K4me3. The numbers of putative enhancers are also shown for each class. Bar chart shows the ratios of top enriched TF motifs (p-value < 1e-20) in putative enhancers that contain CGs for each class (right).

Finally, we asked if ectopically activated enhancers can indeed recruit TFs. The pluripotency factor SoxB1 is critical for ZGA and early development in zebrafish (*74*). SoxB1 includes six Sox genes sox1a/1b/2/3/19a/19b (*75*). We chose Sox2, one of the earliest zygotic genes activated (4.3 hpf) (*76*) for which an antibody is conveniently available, and performed CUT&RUN (*61*) in control and *dnmt1* mKD embryos at shield stage (Fig. S11A). Reassuringly, Sox2 binding in both control and mKD mutants enriches for Sox2 motif (Fig. S11B). Ectopic Sox2 binding occurs in at least 570 ectopic H3K4me3 sites, preferentially aligning at centers of these H3K4me3 peaks (Fig. 6E-F). About 12.7% (186/1,463) of ectopic H3K4me3 sites with *sox2* motif acquire Sox2 binding, while the number decreased to 7.8% (384/4,917) for ectopic H3K4me3 without *sox2* motif (p-value = 3e-8) (Fig. S11C). These Sox2 ectopic binding sites are also closer to derepressed genes in mKD mutants, although it did not reach statistical significance due to the limited numbers of genes (Fig. S11D). Hence, ectopically activated adult enhancers recruit TFs and are correlated with gene derepression.

### Embryonic and adult enhancers exhibit distinct DNA methylation sensitivity and CG densities

While DNA methylation appears to repress adult enhancers, intriguingly, it was reported that embryonic enhancers are hypermethylated and hence are insensitive to DNA methylation in zebrafish early embryos (*39, 41*). This is attributed to the absence of TET proteins, the key regulatory enzymes of DNA demethylation, as its expression is not detectable in zebrafish embryos until 24hrs post fertilization (the phylotypic stage) (Fig. S11E) (*40*) (*39*). It remains elusive why DNA methylation can repress adult enhancers but not embryonic enhancers. Furthermore, given the large numbers of adult enhancers present in the genome (n=60,728 across 11 adult tissues) (Fig. 6G), clearly not all adult enhancers are derepressed and acquire ectopic H3K4me3 in *dnmt1* mutant embryos. To understand why certain enhancers are selectively sensitive to DNA methylation and are prone to ectopic H3K4me3 acquisition, we identified early embryonic enhancers (dome and shield stages) and adult enhancers (across 11 adult tissues) using distal H3K27ac and removed those overlapping with annotated promoters or H3K4me3 in adult tissues (promoter mark) (Methods). We confirmed that while both carry H3K27ac (as defined), embryonic enhancers and adult enhancers are hypermethylated and hypomethylated, respectively (Fig. 6H and S11F). Interestingly, adult enhancers, but not embryonic enhancers, showed elevated CG densities compared to the background (Fig. 6H). Importantly, such high CG density of adult enhancers is much more evident for those that acquire ectopic H3K4me3 upon the loss of DNA methylation (H3K4me3+), but less so for those that did not gain ectopic H3K4me3 (H3K4me3-) (Fig. 6H). This is consistent with the notion that CG-rich sequences can attract histone methyl-transferases such as MLL1/2, which contain the CXXC domain that recognizes unmethylated CpG regions (*77, 78*). Furthermore, adult enhancers that show ectopic H3K4me3 are generally inaccessible in control early embryos, but become accessible in *dnmt1* mKD mutants (Fig. 6I). By contrast, H3K4me3- adult enhancers are weakly accessible, and such accessibility are even somewhat decreased upon the loss of DNA methylation. Interestingly, such difference appears to be also true for embryonic enhancers. A small portion of embryonic enhancers (7.4%, n=726) also acquired H3K4me3 upon the loss of DNA methylation (Fig. 6I). Despite the overall low CG density of embryonic enhancers, these H3K4me3+ embryonic enhancers also showed a slightly higher CG density compared to H3K4me3- enhancers (Fig. 6I). Importantly, only H3K4me3+, but not H3K4me3-, embryonic enhancers showed increased chromatin accessibility in *dnmt1* mKD embryos (Fig. 6I). Notably, the CG densities of H3K4me3+ embryo-specific enhancers are overall still low compared to adult enhancers (Fig. 6I and S11G). In sum, these data revealed that adult enhancers are preferentially CG-rich, more sensitive to DNA methylation, and are prone to acquire ectopic H3K4me3 and increased chromatin accessibility upon the loss of DNA methylation.

Enhancers are activated by interacting TFs (*69, 79*). Therefore, we asked if the differential activation of H3K4me3+ and H3K4me3- adult enhancers may be related to different sets of TFs and whether these TFs are present in early embryos. By searching for TF motif in these enhancers, we found distinct motifs present between H3K4me3+ and H3K4me3- adult enhancers, as well as between adult active enhancers (H3K27ac+) and adult decommissioned enhancers (H3K27ac- but ATAC+/LMR+) (Fig. S11H). Importantly, most TFs of which the motifs are found in H3K4me3+ adult enhancers are expressed in both embryos and adult tissues. By contrast, many TFs of which the motifs are enriched in H3K4me3- adult enhancers are highly expressed in adult tissues but not in early embryos (Fig. S11H, red arrow). In this analysis, we averaged gene expression for TFs from the same family but with almost identical motifs (such as *gata*, *fox* etc.). Hence, the activation of H3K4me3+ adult enhancers in *dnmt1* mKD embryos may be due to both their sensitivity to DNA methylation and the presence of corresponding TFs in early embryos. Meanwhile, the GREAT analysis (*80*) revealed that genes near H3K4me3+ adult enhancers are enriched for those functioning in FGF and Wnt signaling pathway; on the other hand, H3K4me3- adult enhancers are more enriched for kinase signaling pathway, regulation of cell cycle, etc. (Fig. S11I). Finally, we reasoned that if DNA methylation interferes TF binding at CG-rich enhancers, these TFs may be more likely to contain CG in their motifs. Indeed, TF motifs identified from embryonic enhancers are less likely to contain CGs than those from adult enhancers (Fig. 6I, far right, bar chart). However, exceptions are TF motifs identified in both H3K4me3+ embryonic and adult enhancers, which preferentially possess CGs. In sum, these data suggest that adult enhancers are preferentially CG-rich and interact with CG-containing TFs. By contrast, embryo-specific enhancers tend to be CG-poor and interact with CG-less TFs. Given the absence of Tet proteins in early embryos (Fig. S11E) (*39*), these data suggest that the inherited DNA methylation, coupled by enhancer dememorization, presents an epigenetic gate that prevents premature firing of adult enhancers and transcription programs without interfering embryonic programs (Fig. 7A).

**Fig. 7.**
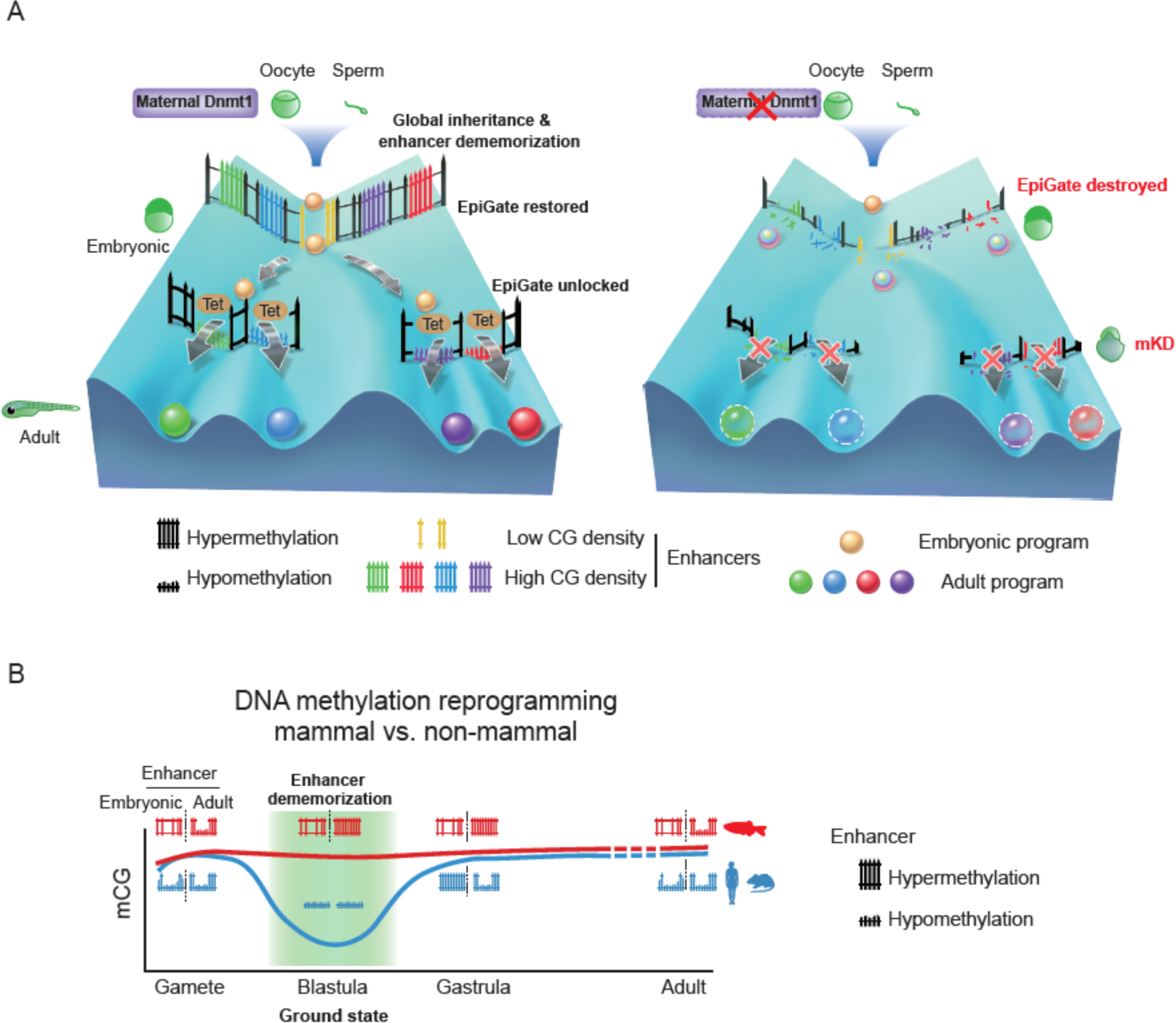
Inherited methylome coupled by enhancer dememorization resets an epigenetic gate that safeguards embryonic programs. **(A)** Maternal Dnmt1mediated inherited global DNA methylome coupled with enhancer dememorization plays an instrumental role in restoring a full methylome to ultimately safeguard embryonic development against premature activation of adult programs. In WT embryos, inherited methylome sets up an “EpiGate” after fertilization and effectively repress adult enhancers, which are preferentially CG-rich. Embryonic enhancers, which are CG-poor, are however insensitive to DNA methylation, and can function while hypermethylated and instruct embryonic transcription program. At the phylotypic stage, expression of *tet* genes demethylate CG-rich adult enhancers, allowing their activation and cell lineage differentiation. While in maternal *dnmt1*mKD embryos, inherited DNA methylome failed to be maintained after fertilization, hence destroying the “EpiGate”. This leads to aberrant activation of adult programs in early embryos, accompanied by developmental failure and embryonic lethality around gastrulation. **(B)** Enhancer dememorization resets the developmental clock by restoring a “ground state” (green shade) free of parental epigenetic memories in both mammals (blue, human and mouse) and non-mammalian vertebrate (red, zebrafish) around blastula stage. Such epigenetic resetting is achieved through global DNA remethylation and demethylation in mammals, and through enhancer hypermethylation in zebrafish. Embryonic enhancers are then demethylated in gastrula only in mammals, where TETs are expressed, but not in zebrafish, where TETs are still silenced. Zebrafish embryonic enhancers are nevertheless functioning presumably due to their low CG densities and insensitivity to DNA methylation. Adult enhancers are subsequently demethylated by TETs in both mammals and zebrafish. In mammals, embryonic enhancers also remain hypomethylated in adult tissues despite being decommissioned (*16, 17*).

## Discussion

The reprogramming of DNA methylation in mammals is critical for successful parental-to-embryonic transition and epigenetic memory resetting between generations. However, many non-mammalian vertebrates appear to lack such global reprogramming (*22–25*). To date, why DNA methylation undergoes such distinct reprogramming modes between mammals and non-mammals remain elusive. This is particular intriguing given histone marks undergo global resetting in both mammals and non- mammalian vertebrates (*26–29, 81*). Here, we sought to decipher this mystery by depleting maternal *dnmt1* in zebrafish early embryos, which revealed an essential role for inherited DNA methylation in early embryonic development. Moreover, such methylome, when coupling with enhancer dememorization, restores a full methylome to guard against premature activation of adult programs through repressing adult enhancers (Fig. 7A). Hence, enhancer dememorization resets the developmental clock by restoring a “ground state” free of parental epigenetic memories. Interestingly, such epigenetic resetting is similarly achieved in mammals but through distinct paths, as global demethylation and remethylation essentially also remove parental epigenetic memories at enhancers (Fig. 7B). Therefore, enhancer dememorization may underlie and potentially unify distinct epigenetic reprogramming modes between mammals and non-mammalian vertebrates.

### Inherited DNA methylation is essential for early development and proper cell differentiation

Single-cell RNA-seq analysis revealed that a subset of developmental genes showed downregulation in *dnmt1* mKD embryos, which are likely indirectly caused by failed differentiation. Differentiation defects are partially attributed to TE-derepression induced immune response and *p53*-mediated apoptosis and cell cycle arrest, as inhibiting *p53* or immune signaling pathways partially rescued the differentiation defects. However, most of these animals still experience embryonic lethality, suggesting that such developmental defects likely stem beyond transposon derepression. In fact, we recently showed that in mouse ESCs that are deficient for all regulatory enzymes of DNA methylation (DNMT3A/3B/3C and TET1/2/3) except for DNMT1, the global DNA methylome is well maintained but becomes static (Wang et al., 2020). While the silencing of transposons is expected to be not affected, this mESC line still failed to differentiate, suggesting that TE-silencing independent function of DNA methylation may also contribute to differentiation defects.

### Promoter DNA methylation reprogramming during the oocyte-to-embryo transition

Despite the global inheritance of methylome, dynamic methylation reprogramming does occur at regulatory elements such as promoters and enhancers in early zebrafish embryos (*24, 25, 29*). In particular, the maternal methylome is conformed to a state that highly resembles that of sperm, including both gain and loss of DNA methylation at specific promoters. Notably, such reprogramming does not depend on sperm, as it can occur even when sperm DNA was disrupted (*25*). Therefore, we previously proposed that both oocyte and sperm are perhaps reprogrammed to an “embryonic state” (*29*). While such reprogramming occurs after fertilization for the maternal genome, it may occur even before fertilization for sperm, as supported by the full methylation of enhancers in sperm. The significance of this intriguing phenomenon, however, remained unclear. One plausible possibility is that such transformation may help prepare gene activation and silencing during the forthcoming ZGA. However, our data showed that among genes that are hypomethylated in oocytes but become hypermethylated in embryos, only a small subset of genes are derepressed in *dnmt1* mKD embryos. The majority of these genes remain silenced by shield stage (Fig. S9C-D), suggesting that such conversion do not seem to solely serve the immediate gene repression after fertilization. Alternatively, such reprogramming may be important not only for ZGA, but also for future development. By resetting adult and gametic epigenetic memories to a “ground state”, DNA methylation reprogramming may facilitate future gene regulation when stage-specific activators and repressors of promoters and enhancers appear in a spatiotemporally controlled manner.

### Reprogramming of enhancers resets an epigenetic gate that prevents precocious activation of adult programs

The idea that DNA methylome reprogramming may have a larger impact beyond ZGA and create a ground state is strongly supported by further analyses of enhancers. Nearly all enhancers in zebrafish gametes are dememorized through DNA hypermethylation in early development (*29*). This essentially creates a methylome free of past enhancer memories and thus represents a likely “ground” epigenetic state. Moreover, hypermethylation of enhancers also prevents precocious activation of adult enhancers in early embryos.

Mechanistically, this is probably due to the absence of Tet proteins which are not expressed until the phylotypic stage. By contrast, mammalian TET proteins are expressed throughout pre- and post-implantation stages and play critical roles at enhancers during gastrulation (*9, 82*). The motivation underlying Tet’s absence in early zebrafish embryos remains unknown. One possibility may lie in the different cell cycle speed in early embryos between mammals (such as 24hrs per cell cycle for cleavage-stage mouse embryos) and cold-blooded vertebrate animals (such as 15 min per cell cycle for zebrafish pre-ZGA embryos). It is tempting to speculate that TETs, even if expressed, may not properly function at enhancers in such rapidly dividing cells. After 24hrs post fertilization, the cell cycle prolongs to more than 3hrs (*32*), which is perhaps more accessible for the epigenome editing enzymes such as Tets.

The absence of Tets presumably also creates a potential challenge for embryonic enhancers to function given their hypermethylated states. Interestingly, our analyses revealed that embryonic enhancers tend to be CG-poor, and are thus less likely to be affected by DNA methylation. In addition, TFs that potentially bind these enhancers tend to contain fewer CGs in their recognition motifs (Fig. S11H). We propose that these enhancers may have adopted these sequence features during evolution to survive without TETs. By contrast, adult enhancers tend to be relatively CG rich and are bound by TFs that contain CGs in their motifs. Upon the loss of DNA methylation, many putative CG-rich adult enhancers become aberrantly activated, as indicated by their acquisition of active marks such as H3K4me1, H3K27ac, H3K4me3, increased chromatin accessibility, and precocious activation of their neighbor genes. Therefore, these data demonstrate that DNA methylome may play a critical role in ensuring temporally ordered enhancer activation during development. Collectively, our data showed that the inherited global DNA methylome, coupled by enhancer dememorization, plays an essential role in embryonic development by repressing transposons and restoring an epigenetic gate that guards against premature activation of adult programs (Fig. 7A). Future studies are warranted to determine whether similar mechanisms (such as enhancer dememorization) can be applied to other reprogramming processes to reset the epigenetic clock and restore cells from differentiated or aged states back to a ground state of totipotency.

## Materials and Methods

### Zebrafish strain and fertilized egg microinjection

The wild type AB (female) strains were used in most experiments. The *p53 ^M214K^* line (*51*) and the *dnmt1^s872^* (*31*) lines were described previously. Embryos derived from *dnmt1^s872^* heterozygous intercrosses were identified by PCR genotyping at desired stages. Ethical approval was obtained from the Animal Care and Use Committee of Tsinghua University. All experimental animal procedures were performed under anesthesia, and all efforts were made to minimize suffering.

For 1-cell microinjection, mouse Stella mRNAs were injected into naturally fertilized 1-cell stage embryos, according to a commonly used zebrafish microinjection protocol (*83*). After injection, embryos were grown in fresh Holtfreter solution (0.05 g/L KCl, 0.1 g/L CaCl2, 0.025 g/L NaHCO3, 3.5 g/L NaCl, pH 7.0) at 28.5℃ and were staged according to standard morphological criteria (*32*). The dose of mouse Stella mRNA was 550 pg per embryo.

### *dnmt1* mKD with oocyte microinjection *in situ* (OMIS)

Briefly, on the first day of OMIS (*42*), adult females at 5-12 month old were anesthetized in 550 µg/ml tricaine (Sigma, Cat A5040) in a petri dish. Then the fish was placed on a damp sponge with specific buffer (5.4 mM KCl, 136.8 mM NaCl, 4.2 mM NaHCO3, 0.44 mM KH2PO4, 0.25 mM Na2HPO4, and 0.5% (wt/vol) BSA). A cut was made on one side of belly to expose the ovary. The diluted MOs were microinjected into each oocyte. Rhodamine B (Sigma, Cat R8881) was co-injected with MOs as a dye. After injection, the wound on the belly was sewed with a surgical sewing needle carefully and quickly. Once the operation was done, the female was transferred into fish water supplemented with 32 µg/ml tricaine, 10 unit/ml penicillin and 10 µg/ml streptomycin (HyClone, Cat SV30010). Then the fish was transferred to fish water containing gradually reduced concentrations of tricaine. In the evening of the second day, the female was paired with a wild type male. In the morning of the third day, the pair started to chase and lay fertilized eggs naturally. The injected oocyte-derived embryos were identified by co-injected dye (rhodamine B) at very early developmental stages. The injection doses of *dnmt1*-MO and standard control MO (cMO) were both 5 ng per oocyte in the same assay. The sequences of MOs are 5’- ACAATGAGGTCTTGGTAGGCATTTC-3’ (*dnmt1*-MO) (*84*), and 5’-CCTCTTACCTCAGTTACAATTTATA-3’ (cMO). MOs were dissolved in RNase-free water, and heated to 65°C for 10 min before microinjection.

### Tissue collection

Embryos at 5 days post fertilization were euthanized with tricaine, and head and tail were dissected carefully by tweezers. After brief grinding, tissues were frozen at -80℃ for later usage.

### Immunofluorescence and imaging

The embryos at the defined time points after fertilization were fixed by 4% polyformaldehyde overnight at 4℃. Then they were dechorionated manually and dehydrated with methanol. The whole-mount immunofluorescences with Dnmt1 antibody (Santa Cruz, Cat sc-20701), 5mC antibody (Abcam, Cat ab10805), and pH2A.X antibody (Cell Signaling, Cat 2577S), were done with DAPI (Invitrogen, Cat D1306) staining and performed as previously described (*85*). The secondary antibodies were Alexa Fluor® 488 conjugated anti-rabbit and Alexa Fluor® 488 conjugated anti-mouse (Jackson ImmunoResearch, 1:200 diluted). After staining, embryos were deyolked by tweezers and mounted on glass slides in mounting medium (Sigma, Cat P3130) at animal polar upturned position. Images were acquired on 710 or 880 META laser scanning confocal microscope and manipulated by ZEN software.

Treated or untreated embryos were anesthetized at desired stages with 0.02% tricaine and mounted in 5% methyl cellulose (Sigma, Cat M-6385) for observation, and phenotype pictures were taken under Nikon SMZ1500 microscope.

### *dnmt1* rescue

The full-length *dnmt1* coding sequence from zebrafish was cloned into pXT7 vector and linearized by SmaI digestion. mRNA was synthesized in vitro using mMESSAGE mMACHINE kit (Ambion, Cat AM1344) and purified using RNeasy Mini kit (Qiagen, Cat 74104). To avoid binding of *dnmt1* mRNA with MO, the target sequence of MO was mutated without affecting amino acid sequence (WT mis-*dnmt1* mRNA). Catalytic mutant *dnmt1* mRNA was designed according to *s872* mutant (*31*), which contains a stop codon in catalytic domain resulting the loss of function of Dnmt1 (Mut mis-*dnmt1* mRNA). These mRNAs were co-injected with MO into GV oocytes by OMIS (*42*) for rescue experiment. Embryos at desired stages were then collected for further analysis.

### Mouse Stella overexpression

The construct containing full-length Stella coding sequence from mouse was generated and linearized by NotI digestion. The mutant Stella (KRR, K85E/R86E/R87E) contains 3 amino acids mutations within the nuclear export signal as previously described (*35*). mRNA was synthesized in vitro using mMESSAGE mMACHINE kit (Ambion, Cat AM1344) and purified using RNeasy Mini kit (Qiagen, Cat 74104). These mRNAs were injected into zygotes, and embryos at desired stages were collected for further analysis.

### TUNEL

ApopTag Red In Situ Apoptosis Detection Kit (Millipore, Cat S7165) was used to probe cell apoptosis in zebrafish embryo as previously described (*86*). Collected embryos at desired stages were fixed by 4% polyformaldehyde overnight at 4℃, then dechorionated manually and dehydrated with methanol. After being stored at -20℃ for 1hr, embryos were rehydrated with 0.1% PBST (0.1% Triton-X 100 in PBS), then re-fixed with 4% polyformaldehyde for 20min at room temperature and put into precooled ethanol-ethyl acetate mixture (volume ratio 2:1). Next, 50 µl equilibration buffer was added into tubes contained embryos at room temperature for 1hr, and changed with 55 µl TdT reaction system (38.5 µl reaction buffer and 16.5 µl TdT enzyme) at 37℃ more than 1hr to add Digoxin labeled dUTP in DNA breaks. To stop reaction and visualize Degoxin signals, embryos were washed with STOP/Wash buffer and incubated with anti-Digoxin antibody coupled with Rhodamine buffer (34 µl blocking buffer and 31 µl antibody) at 37℃ for 30min or overnight. DNA was stained with DAPI. After staining, embryos were mounted in the same way as immunofluorescence and imaged on 710 or 880 META laser scanning confocal microscope.

### Inhibitor treatment

To inhibit the reverse transcription, embryos were treated with 50 μM Foscarnet (Selleck, Cat S3076). To inhibit the immune response, embryos were treated with 0.1 μM or 0.01 μM BX795 (Selleck, Cat S1274). Embryos treated with DMSO were used as control. All these embryos were examined for phenotypes and fixed for immunostaining at desired stages. Phenotype at 24hrs post fertilization was quantified and summarized.

### STEM-seq library preparation

STEM-seq was carried out as described previously (*43*). The deyolked embryos and tissues were lysed with 20 µl lysis buffer (10 mM Tris-HCl, pH 7.4, 10 mM NaCl, 3 mM MgCl2, 0.1 mM EDTA, pH 8.0, 0.5% NP-40) and 2 µl protease K (Roche, Cat 10910000) for at least 3hrs at 55℃. After heat-inactivation, spike-in λ-DNA (Promega, Cat D150A) was added at a mass ratio of 1/200. Bisulfite conversion was performed with the EpiTect Fast Bisulfite Conversion Kit (Qiagen, Cat 59824). The converted DNA was subjected to column purification and desulfonation on MinElute DNA spin columns (Qiagen, Cat 59824) with carrier RNA (Qiagen, Cat 59824) according to the manufacturer’s instructions. The purified DNA was eluted in 30 µl of elution buffer and ready for TELP library preparation (*87*).

### Total RNA-Seq library preparation and sequencing

The embryos were dechorionated manually by tweezers and transferred into 750 µl Trizol (Invitrogen, Cat 15596018). About 10 fresh embryos were transferred into Trizol and vortexed until no visible particles. 150 µl chloroform (Amresco, Cat 0757) was added and mixed thoroughly. The mixture was then transferred into phasemaker tube (Invitrogen, Cat A33248) and spun at 14,000 rpm for 15 min. Next, the top phase was taken out from the tube, and 1 µl LPA was added (Sigma, Cat 56575) and mixed well using pipettes. Then, RNA was precipitated by adding 750 µl isopropanol (Sigma, Cat 59304) at -20℃ overnight. At the next day, the tube was spun at 14,000 rpm for 30 min and supernatant was removed. The pellet was washed with fresh 70% ethanol, re- suspended in 20 µl RNase-free water and stored at -80℃ for later usage.

NEBNext rRNA Depletion Kit (NEB, Cat E6310S) was used to deplete ribosomal RNA according to the manufacture’s instruction. Briefly, rRNA were hybridized with probes and digested with RNase H, then excess probes were digested with DNase I. After that, NEBNext RNA sample purification beads were used to purify rRNA depleted RNA. Purified RNA was fragmented before cDNA synthesis at 95℃ for 8 min. Double-stranded cDNA was synthesized with NEB Next first-strand (NEB, Cat E7771S) and second-strand synthesis modules (NEB, Cat E7550S), then purified with Ampure XP beads (Beckman, Cat A63882). Synthesized cDNA was subjected to library preparation with NEBNext Ultr II DNA Library Prep Kit (NEB, Cat E7645S). DNA was end-repaired, adenylated, and ligated to TruSeq sequencing adaptors. DNA was amplified using KAPA HF HotStart ReadyMix (KAPABiosystem, Cat RR2602). The amplified DNA was size-selected using Ampure XP beads for 200-500 bp DNA fragments. All libraries were sequenced by Illumina Hi-Seq 1500 or 2500 or XTen platform according to manufacturer’s instruction.

### Single-cell RNA-seq library preparation and sequencing

Cell dissociation protocol was based on a previously described method (*48*) with modifications to adapt it for 10× Genomics platform. Briefly, *dnmt1* mKD and control embryos were collected 20 min after fertilization and cultured as mentioned before. Then 15 embryos at dome or shield stages for each sample were transferred into plastic Petri dishes that had previously been coated with 2% agarose and soaked with DMEM/F12 medium (Gibco/Life Technologies, Cat 11330032) at least 2hrs. Next, embryos at desired stages were dechorionated and deyolked manually by forceps and transferred to a 1.5 ml Eppendorf tube with 50 µl DMEM/F12 medium. Dissections were performed for up 15 min. The volume of DMEM/F12 medium containing embryos was adjusted to 200 µl, and then cells were mechanically dissociated by flicking the tube 30 times and pipetting mixture 10 times through a 200 µl tip. The volume was adjusted to 1 ml with PBS containing 1.0% BSA, and spun to pellet cells at 300 g for 30s. The supernatant was removed and cells were resuspended in 80 µl of PBS containing 0.1% BSA and 20% Optiprep (StemCell, Cat 07820), aiming for a concentration above 300 cells/µl. Cells were then passed through a cell sieve (100 µm for dome stage and 70 µm for shield stage).

Single-cell RNA-seq library was performed with Chromium Next GEM Single-cell kit (10× Genomics, Cat PN- 1000121) based on the standard protocol. All libraries were sequenced by Illumina Hi-Seq 1500 or 2500 or XTen platform according to manufacturer’s instruction.

### CUT&RUN library preparation and sequencing

CUT&RUN was conducted as previously described (*61, 88*) with modifications in cell permeation to adapt it for zebrafish embryos. Embryos were deyolked by tweezers manually and transferred into 1.5 mL conventional, non- low-binding tube (Axygen). Then tubes were flicked several times to disperse cells in embryos, and resuspended by 60 ul washing buffer (20 mM HEPES-KOH, pH = 7.5; 150 mM NaCl; 0.5 mM Spermidine and Roche complete protease inhibitor). 10 µl Concanavalin-coated magnetic beads (Polyscience, Cat 86057) for each sample were gently washed twice, resuspended by binding buffer (20 mM HEPES-KOH, pH = 7.5; 10 mM KCl; 1 mM CaCl2; 1 mM MnCl2), and added carefully to the cells. The cells with beads were incubated at 23℃ for 30 min on Thermomixer (Eppendorf) at 400 rpm, then held at magnetic stand to exclude buffer, and resuspended by 75 µl antibody buffer (washing buffer supplied with 0.02% digitonin and 2 mM EDTA, freshly made) with antibodies against H3K4me3 (in-house) (*89*), H3K27me3 (Active Motif, Cat 61017), H2AK119ub (CST, Cat 8240s), or Sox2 (Active Motif, Cat 39843) diluted at ratio of 1:100. Then the samples were incubated at 4℃ on Thermomixer overnight at 400 rpm. On the second day, the samples were washed by digitonin-washing buffer several times on magnetic stand, and resuspended with 50 µl digitonin-washing buffer supplied with 700 ng/ml pA-MNase, and incubated at 4℃ on Thermomixer for 3hrs at 400 rpm. After that, the cells were washed by digitonin-washing buffer on magnetic stand, and resuspended by 100 µl digitonin-washing buffer on ice for at least 2 min. Targeted region digestion was activated by adding 2 µl 100 mM CaCl2 for 30 min in ice, then stopped by 100 µl 2 X stop buffer (340 mM NaCl; 20 mM EDTA, pH = 8.0; 4 mM EGTA, pH = 8.0; 50 µg/ml RNase; 100 µg/ml glycogen; 0.02% digitonin suppled with spike-in DNA) and fully vortexed. To release fragments, the samples were incubated at 37℃ on Thermomixer at 400 rpm for 20 min. Then the supernatants were purified by phenol chloroform and ethanol purification, and subjected to Tru-seq library construction using NEBNext Ultra II DNA Library Prep kit (NEB, Cat E7645S) as standard protocols. The amplified DNA was size-selected using AMPure Beads for 200-800 bp DNA fragments. All libraries were sequenced by Illumina Hi-Seq 1500 or 2500 or XTen platform according to manufacturer’s instruction.

### STAR ChIP-seq library preparation and sequencing

The STAR ChIP-seq was performed as previously described (*29*). Embryos were deyolked by repeatedly blowing with a 200 µl pipette and cell pellets were collected by spinning down at 5000 rpm for 5 min at 4℃. After centrifuge, cell pellet was lysed in 40 µl lysis buffer (0.5% NP-40, 0.5% Tween, 0.1% SDS and proteinase inhibitor) with pipetting up and down several times. 0.1 unit of MNase (Sigma, Cat N3755) was added for chromatin digestion at 37℃ for 5 min. The reaction was then terminated by adding 1 µl 0.5 M EGTA. IP sample was incubated with 1 µg H3K4me3 antibody (in-house) (*89*) and 2 µg H3K27me3 antibody (Active Motif, Cat 61017) overnight with rotation at 4℃. In the next day, the sample was incubated with 300 µg protein A or protein G dynabeads (Life technologies, Cat 10001D or 10003D) for 2hrs with rotation at 4℃. The beads were washed 5 times in 150 µl RIPA buffer and once in 150 µl LiCl buffer. After washing, tubes were spun briefly and the supernatant was removed. For each IP sample, beads were resuspended with 27 µl ddH2O and 1 µl 10× Ex-Taq buffer (TaKaRa). 1 µl proteinase K (Roche, Cat 10910000) was then added and the mix was incubated at 55℃ for 90 min to elute DNA from beads.

The supernatant was then transferred to a new tube and the proteinase K was inactivated at 72℃ for 40 min. 1 µl rSAP (NEB, Cat M0371) was then added to dephosphorylate 3’ end of DNA at 37℃ for 1hr. rSAP was inactivated at 65℃ for 10 min. The resulting sample was subjected to TELP library preparation as previously described (*87*). The amplified DNA was size-selected using AMPure Beads for 200-800 bp DNA fragments. All libraries were sequenced by Illumina Hi-Seq 1500 or 2500 or XTen platform according to manufacturer’s instruction.

### ATAC-seq library preparation and sequencing

The miniATAC-seq procedure was performed as previously described (*71*) with modifications to adapt it for zebrafish embryos. Briefly, dispersed cells from deyolked zebrafish embryos were transferred into 6 µl lysis buffer (10 mM Tris-HCl, pH = 7.4; 10 mM NaCl; 3 mM MgCl2; and 0.02% digitonin) in ice for 10 min. The ATAC reaction was performed by adding 4 µl ddH2O, 4 µl 5 X TTBL, and 5 µl TTE mix V5 (Vazyme, Cat TD502) at 37℃ for 30 min, then stopped by adding 5 µl 5 X TS stop buffer at room temperature for 5 min. DNA was extracted by phenol chloroform and ethanol purification after adding 40 ng carrier RNA and 103 µl Tris-EDTA. Then DNA was PCR amplified with 10 µl index (Vayzme, Cat TD202), 10 µl 5 X TAB and 1 µl TAE (Vayzme, Cat TD502), with the program of 72℃ for 3 min, 98℃ for 30s, (98℃ for 15s, 60℃ for 30s, and 72℃ for 3min) with 18 cycles, and 72℃ for 5 min. The amplified DNA was size-selected using AMPure Beads for 200-800 bp DNA fragments. All libraries were sequenced by Illumina Hi-Seq 1500 or 2500 or XTen platform according to manufacturer’s instruction.

### STEM-seq data processing

All STEM-seq datasets were mapped to the danRer7 reference genome by Bismark (*90*). Reads were trimmed with cutadaptor (*91*) using parameters: --minimum-length 20 --pair-filter=any. Alignments were performed with the following parameters: -N 1 -X 600 --score_min L,0,-0.6. Multi-mapped reads and PCR duplicates were removed. The function bismark_methylation_extractor was used to calculate the DNA methylation level.

### Total RNA-seq data processing

The total RNA-seq data were firstly processed using Trim Galore! with default parameters to trim the adapter- containing and low-quality reads. The filtered data were then mapped to the zebrafish reference genome (danRer7) by STAR (version: STAR_2.5.3a_modified) (*92*) with the following parameters: --outSAMstrandField intronMotif --outSAMattributes All --outSAMunmapped Within --outSAMattrIHstart 0 --outWigStrand Stranded -- outFilterMultimapNmax 20 --twopassMode Basic. The gene expression level was normalized to fragments per kilobase of transcript per million mapped (FPKM) values using Cufflinks (version 2.2.1) (*93*).

### Single-cell RNA-seq data processing

The scRNA-seq data were processed with Cell Ranger 3.1 for genome alignment (danRer10) and transcript counting. Next, the raw count matrices were imported to Seurat 3.0 for identification of cell subtypes. To filter low-quality cells, cells with unique detected genes lower than 2,000 and higher than 6,000 genes were discarded, and cells with high percentages of mitochondrial genes (>5%) were removed too. To make data comparable between cells and samples, we performed SCTransform normalization for each dataset independently. To further eliminate the effects caused by sequencing depths and percentages of mitochondrial genes, UMI variance and percent.mt were regressed out from SCTransform normalized data with regularized negative binomial model and linear model respectively. Samples of dome and shield stages were integrated by identifying anchors with the top 3,000 gene features. During the integration, the control cells were set as reference. Dimensional reduction was performed on the integrated data with PCA and then the first 30 PCs were selected for cluster identification and UMAP visualization.

Positive marker genes for each single cell subtype were identified as following: the expression of a gene is detected in at least 25% of cells of current cell subtype, and its expression is higher than the average expression levels of the rest cells (average log (fold change)>0.25, FDR<0.01).

### Cellular trajectory analysis

To better study the major differentiation lineages, we removed identified apoptotic cells, EVL, YSL, and PGC cells which are more distally related lineages. Raw count matrices were imported to Monocle 2 and the lists of differentially expressed genes in each cluster identified through Seurat were used for the construction of cell trajectory. The root of each trajectory was defined by epiblast cells.

### CUT&RUN, ChIP-seq and ATAC-seq data processing

All reads were aligned to the zebrafish reference genome (danRer7) using Bowtie2 (version 2.2.2) (*94*) with the parameters –t –q –N 1 –L 25. All unmapped reads, non-uniquely mapped reads and PCR duplicates were removed. For downstream analysis, the read counts were normalized by computing the numbers of reads per kilobase of bin per million of reads sequenced (RPKM). To minimize the batch and cell type variation, RPKM values across whole genome were further Z-score normalized. To visualize the signals in the UCSC genome browser, each read was extended by 250 bp and the coverage for each base was counted.

### Differential gene expression analysis

Read counts of genes were summarized with featureCounts. Next, the read count matrices were imported to DESeq2 to perform differential expression analysis for genes in control and *dnmt1* mKD embryos. Genes with FDR<0.01 and log2(fold change)>2 were considered as significantly differentially expressed genes.

### Analysis of maternal, dome-specific and shield-specific genes

The maternal genes were identified by high expression level in oocyte (FPKM>10), but low at post-ZGA stages (fold change of dome/oocyte<0.5). Dome-specific expressed genes were defined by low expression in oocyte and the 256-cell stage (FPKM<5), and high expression at dome stage (FPKM>10). Similarly, shield-specific genes were defined with low expression levels in oocyte, embryos at the 256-cell and dome stages (FPKM<5), and high expression at shield stage (FPKM>10).

### Analysis of non-maternal ZGA genes

Due to the possible confounding effects from maternally inherited RNA transcripts, non-maternal ZGA genes were defined as low expression level in oocyte and the 256-cell stage (FPKM < 5), but high expression level at dome stage (FPKM of dome>5 and fold change of dome/256-cell > 2).

### Gene ontology analysis

Functional annotation was performed using the Database for Annotation, Visualization and Integrated Discovery (DAVID) bioinformatics resource (*95*). Gene ontology terms for each functional cluster were summarized to a representative term, and P values were plotted to show the significance.

### Identification of CUT&RUN, ChIP-seq and ATAC-seq peaks

Peaks were called using MACS2 (*96*) with the parameters -p 1e-5 --nomodel -g 1.3e10. The called peaks with weak signals were filtered in the further analysis. Where appropriate, a random set of peaks with identical lengths were used as controls.

### Identification of putative enhancers

For adult enhancers, H3K27ac data were used to identify active enhancers, and ATAC-seq and DNA methylation data were used to identify decommissioned enhancers (*17*). In order to ensure the consistency of data processing, the H3K27ac, ATAC-seq and DNA methylation data collected from previous study (*56*) were reanalyzed with our pipelines as described above. The lowly methylated regions (LMRs) were obtained from the same study (*56*).

To define active enhancers, H3K27ac peaks at least +/- 2 kb away from annotated promoters were firstly selected as candidate enhancers. To exclude possible unannotated promoters, any H3K27ac sites that overlap with H3K4me3 peaks of adult tissues (*56*) were also excluded. Furthermore, to exclude possible unannotated promoters in embryos, we took advantage of the notion that hypomethylated regions often mark promoters in zebrafish (*97*) and enhancers are hypermethylated in zebrafish early embryos, leaving only promoters unmethylated (*29*). Therefore, we chose the 256-cell stage as a representative stage when enhancer dememorization is completed. All UMR/LMRs (*29*) at the 256-cell stage of WT embryos are merged together and identified as possible promoters, and were further excluded from the enhancer lists to generate the final putative enhancer lists.

### TF motif identification at enhancers

The findMotifsGenome.pl script from HOMER program (*98*) was used to identify the enriched motifs at enhancers. TFs expressed at least at one stage with motif enrichment *p*-value less than 1e-20 were included.

### Identification of lost, ectopic and retained histone mark peaks

We clustered H3K4me3, H3K27me3 or H2AK119ub peaks and categorized them into lost, ectopic and retained groups based on the following cut-offs. The lost peaks were identified with log2(fold change) <-2, normalized RPKM<-1 in mKD embryonic cells; the ectopic peaks were defined by log2(fold change)>2, normalized RPKM>1 in mKD embryonic cells; the retained peaks were defined by log2(fold change) <2, normalized RPKM>0.5 in both control and mKD samples.

### Distance calculation for ectopic distal H3K4me3 peaks and DEGs

We calculated the distances from the TSSs of upregulated, downregulated or non-differentially expressed genes (non-DEGs) to the center of distal ectopic H3K4me3 peaks. Statistical significance between groups was estimated with Mann Whitney U Test.

### Quantification and Statistical analysis

At least two biological replicates were used for RNA-seq, scRNA-seq, CUT&RUN, STAR-ChIP, and ATAC-seq experiments. The reproducibility between replicates were estimated with Pearson correlation. All box plots were plotted using R and Python. In box plots, the horizontal line shows the median, the box encompasses the interquartile range, and whiskers extend to 5th and 95th percentiles. Statistical significance for the enrichment of *dnmt1* mKD embryos upregulated genes in adult tissue expressed genes was assessed with one-sided Fisher’s exact test.

## Acknowledgments

We appreciate comments from members of the Xie lab during preparation of the manuscript. We would like to express our gratitude to Dr. Sebastian Bultmann and Heinrich Leonhardt from Ludwig Maximilians University of Munich for sharing mouse Stella plasmid and discussion, Dr. Steven Henikoff from Fred Hutchinson Cancer Research Center for sharing pA-MNase and Dr. Qinghua Tao from Tsinghua University for providing the DNMT1 antibody. We are also grateful to the cell facility at the Tsinghua Center of Biomedical Analysis for assistance with imaging and the biocomputing facility at Tsinghua University.

## Funding

National Key R&D Program of China 2016YFC0900300 National Basic Research Program of China 2015CB856201

National Natural Science Foundation of China 31422031 and 31725018 THU-PKU Center for Life Sciences

W.V. is a Howard Hughes Medical Institute international research scholar

## Author contributions

W.W. collected zebrafish samples and conducted most zebrafish experiments with help from W.S. X.W. performed total RNA-seq, scRNA-seq, ATAC-seq, CUT&RUN experiments with the help from L. L. and W. Xia. and performed data analysis with the help from B. Z., H. Z., B. L. B. Z. performed STAR ChIP-seq and STEM-seq experiments with the help from Y. Z. and data analysis. H.Z. performed bulk RNA-seq and scRNA-seq data analyses.

Q.W. helped with mouse Stella overexpression experiments. Xi. W. helped with data analysis. W. Xie. supervised the project or related experiments. X.W., B.Z., H.Z., and W. Xie. wrote the manuscript with help from all authors.

## Data and materials availability

All data have been deposited to the Gene Expression Omnibus (GEO), and the accession number is GSE175951.

