## Supplementary figures for "Methylome inheritance and enhancer dememorization reset an epigenetic gate safeguarding embryonic programs"

**This PDF file includes:**

Figs. S1 to S11

Supplementary Figure 1

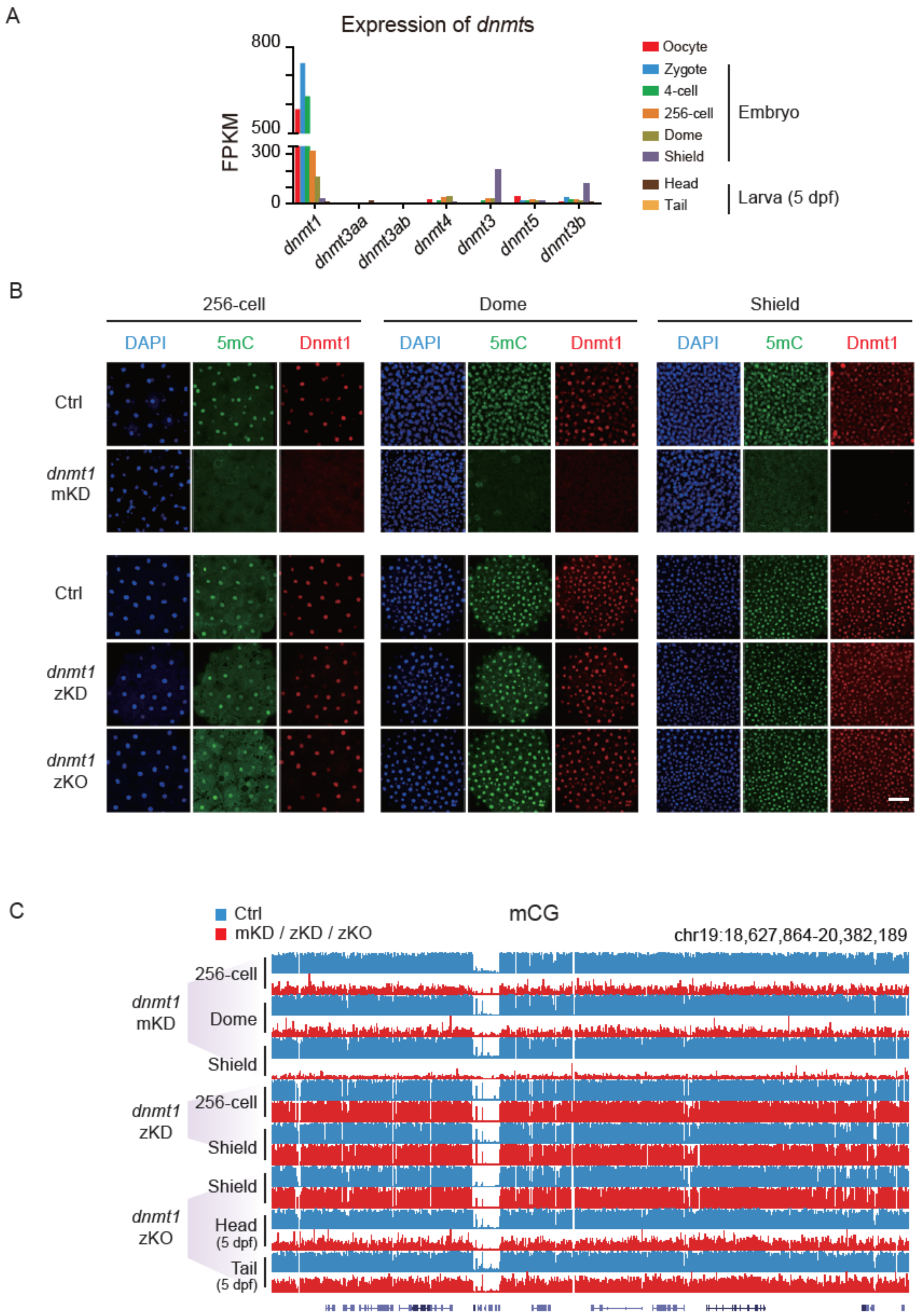


**Fig. S1. Depleting maternal, but not zygotic, *dnmt1* disrupts DNA methylation in early zebrafish embryo.**

**(A)** Expression of zebrafish DNA methyltransferases at various developmental stages. Dpf, days post fertilization.

**(B)** Immunostaining of 5mC (green) and Dnmt1 (red) in control and *dnmt1* mKD, zKD, and zKO embryos at 256-cell, dome and shield stages. DNA was stained with DAPI (blue). Scale bar, 50 μm. **(C)** A UCSC genome browser snapshot showing DNA methylation in control and *dnmt1* mKD, zKD, and zKO embryos at indicated stages.

Supplementary Figure 2

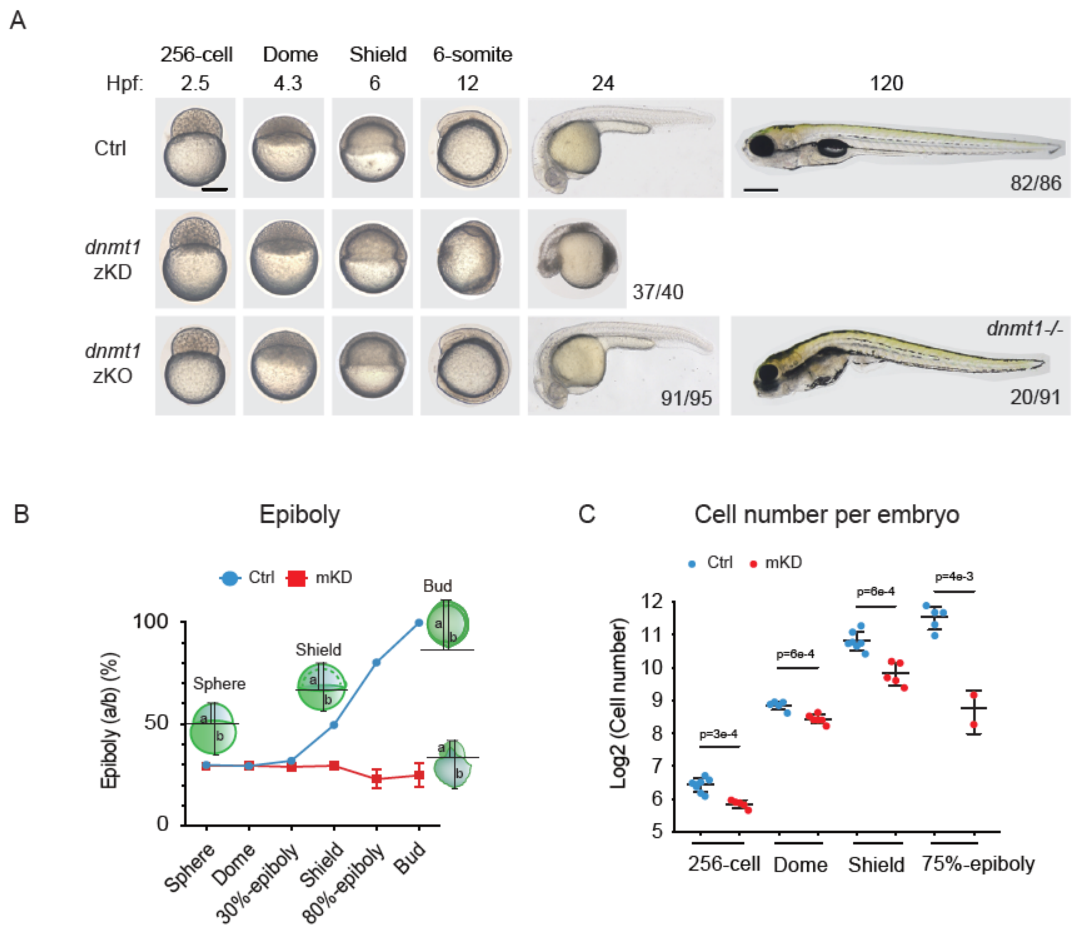


**Fig. S2. Maternal dnmt1 knockdown causes early developmental defects and lethality.**

**(A)** Representative images of embryo phenotypes in control (top), zKD (middle), and zKO embryos (bottom) at 256-cell, dome, shield, 6-somite, 24hpf and 120hpf stages. Scale bar, 250 µm. Hpf, hours post fertilization.

**(B)** Epiboly percentage curve showing the fraction of the yolk cell that is covered by the blastoderm (ratio of a to b) of control (blue) and mKD (red) embryos at sphere, dome, 30%-epiboly, shield, 80%-epiboly and bud stages. Schematic of epiboly percentage (a/b) in control embryos at sphere, shield, bud stages, and mKD embryos at bud stage are shown.

**(C)** Cell number per embryo is shown (based on DAPI staining) in control (blue) and mKD (red) embryos at 256-cell, dome, shield and 75%-epiboly stages.

Supplementary Figure 3

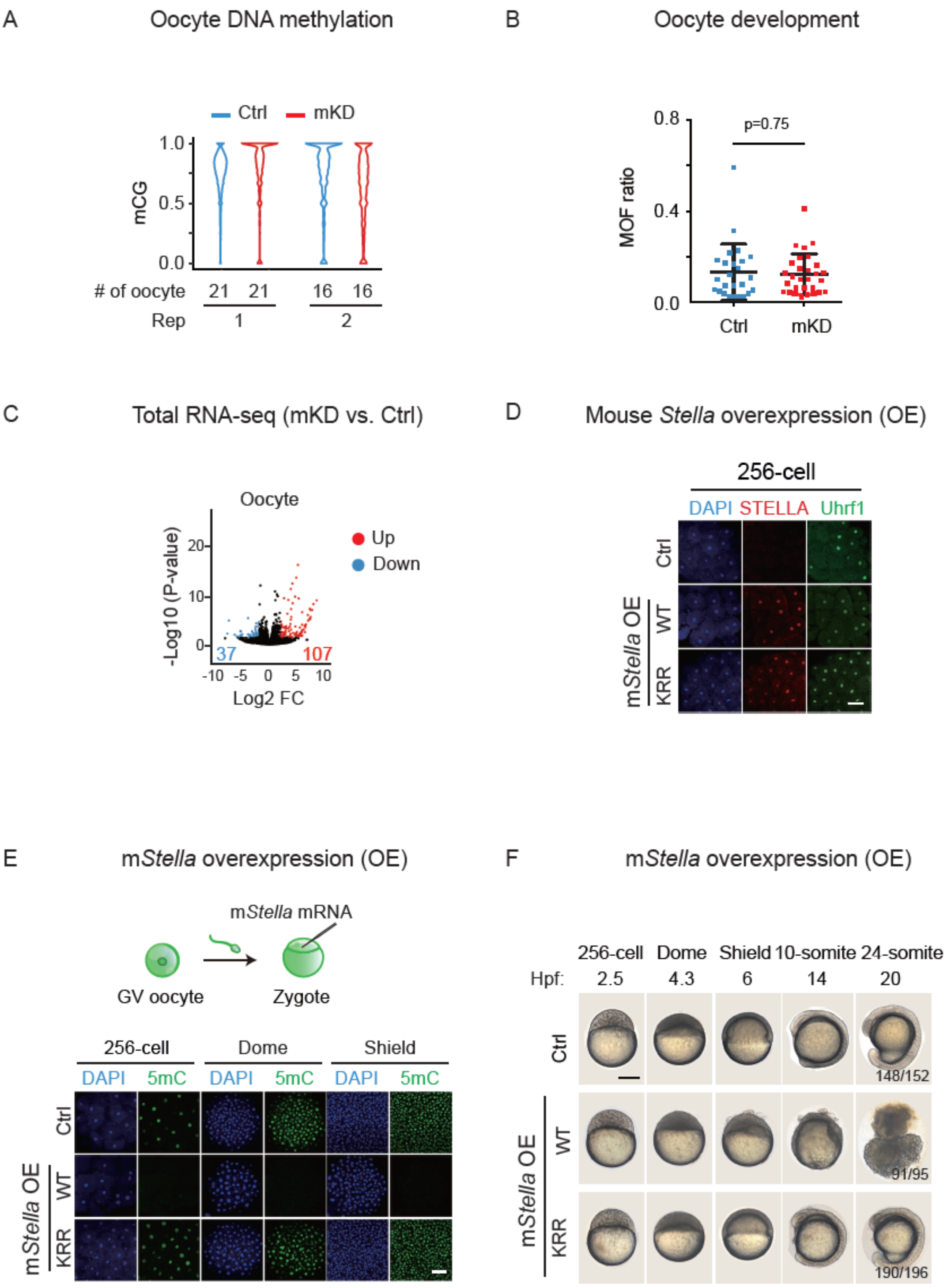


**Fig. S3. Oocyte development was not significantly affected upon *dnmt1* mKD.**

**(A)** Violin plots showing DNA methylation levels across the genome in control (blue) and *dnmt1* mKD (red) oocytes (two replicates). The numbers of oocyte used in each sample are shown below. **(B)** MOF rates (the ratio of maturation, ovulation, and fertilization) of control (blue) and mKD (red) oocytes. Each dot indicates MOF of one injected female with control MO (Ctrl, blue) or *dnmt1* MO

(mKD, red). The numbers of females used in control and mKD groups are 25 and 29, respectively.
**(C)** Volcano plots showing gene expression change (log2(FPKM + 1)) between control and *dnmt1* mKD oocytes. Red dots indicate upregulated genes; blue dots indicate downregulated genes. The numbers of dysregulated genes are also shown.
**(D)** Immunostaining of overexpressed mouse STELLA (mSTELLA) and zebrafish Uhrf1 in zebrafish 256-cell embryos with WT and mutant (KRR, Methods) mouse *Stella* overexpressed. DNA was stained with DAPI (blue). Note STELLA can sequester UHRF1 regardless whether it is in nucleus or cytoplasm (Du et al., 2019; Li et al., 2018; Mulholland et al., 2020). Scale bar, 50 μm.
**(E)** Schematic of mouse *Stella* overexpression in zebrafish zygote and 5mC (green) immunostaining of control and WT or KRR mutant *Stella* overexpressed embryos at different stages. DNA was stained with DAPI (blue). Scale bar, 50 μm.
**(F)** Representative images of uninjected control, and WT or KRR mouse *Stella* overexpressed embryos across different developmental stages. Hpf, hours post fertilization. The numbers of embryos and ratios with a particular phenotype are also shown. Scale bar, 250 μm.

### Supplementary Figure 4

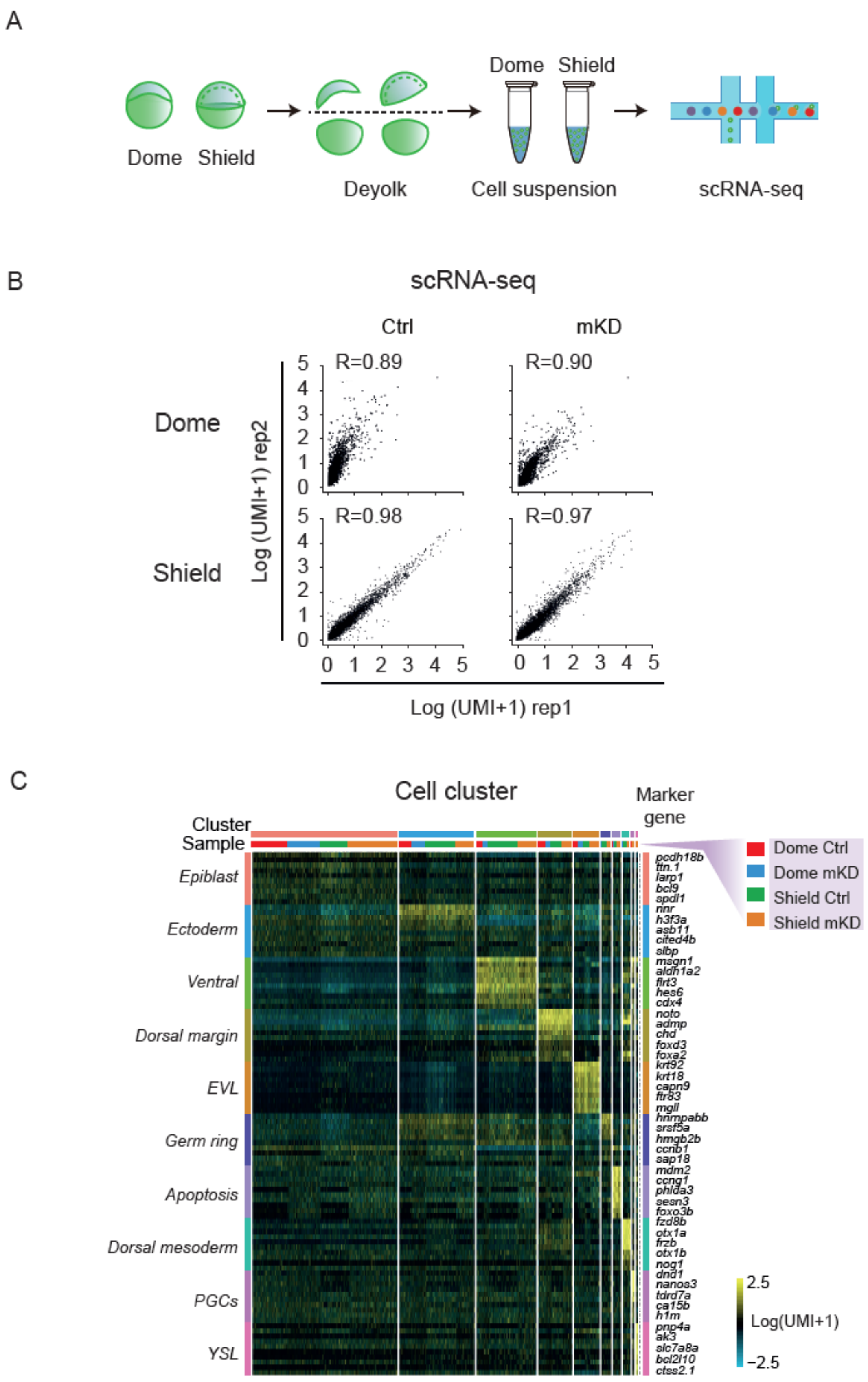

**Fig. S4. scRNA-seq analyses of control and *dnmt1* mKD embryos.**

**(A)** Schematic of scRNA-seq of control and *dnmt1* mKD embryos at dome and shield stages via 10× Genomics platform.

**(B)** Scatter plots comparing gene expression log2 (UMI + 1) between two replicates of control (blue) and *dnmt1* mKD (red) embryos at dome and shield stages.

**(C)** Heat map of top10 marker genes in each cluster in dome control (red), dome mKD (blue), shield control (green) and shield mKD (orange) embryos at shield stage. Cell types are listed on the left, and the top 5 marker genes are listed on the right.

Supplementary Figure 5

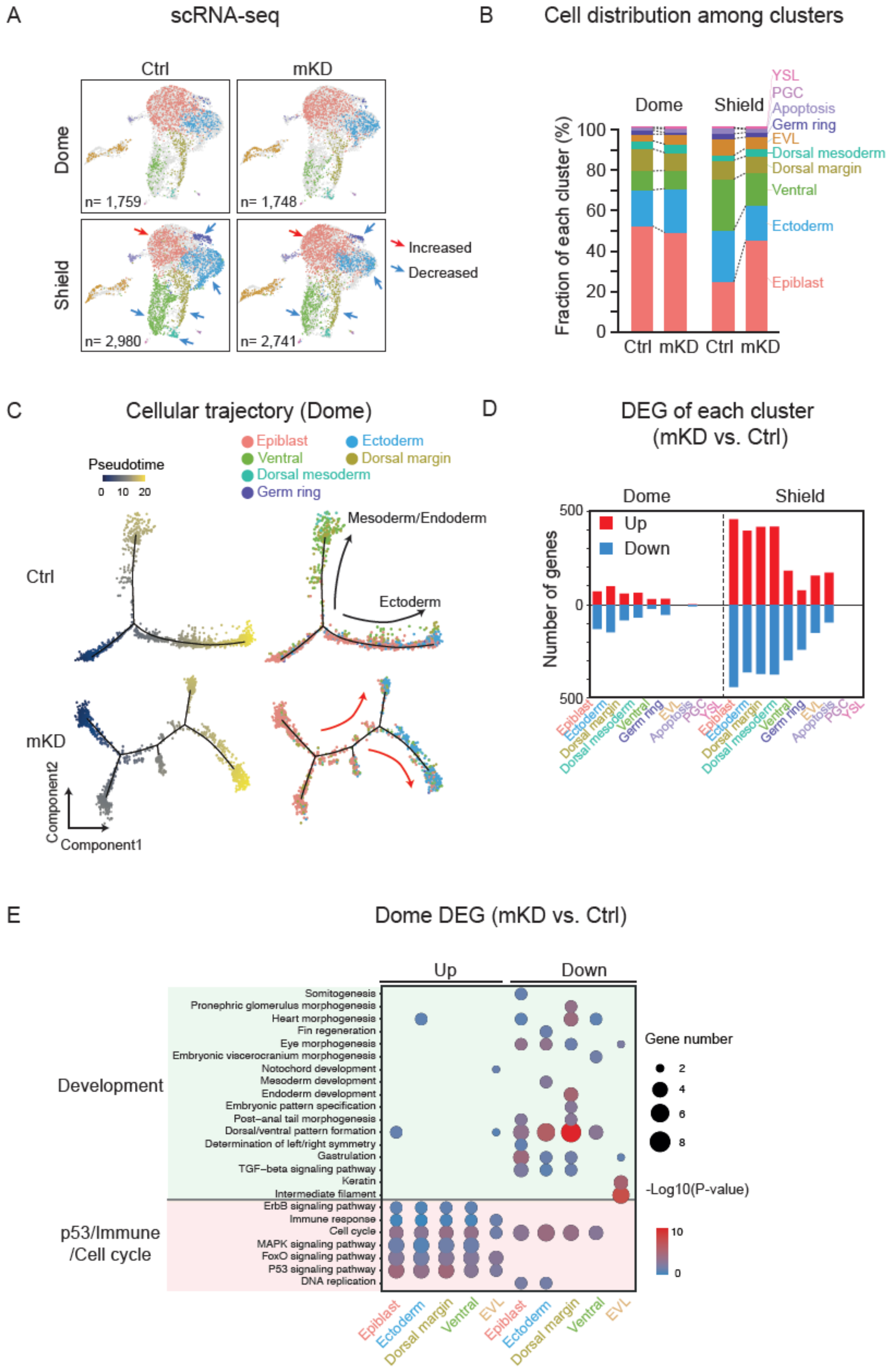

**Fig. S5. scRNA-seq reveals severe differentiation defects in *dnmt1* mKD embryos.**

**(A)** Projection of cells with UMAP for control and *dnmt1* mKD embryos at dome and shield stages. Red arrow indicates a cluster with increased cell numbers in mKD embryos at shield stage. Blue arrows indicate clusters with decreased cell numbers in mKD embryos at shield stage.

**(B)** Bar chart showing cell fraction of each cluster in control and *dnmt1* mKD embryos.

**(C)** Pseudotime trajectory of control and *dnmt1* mKD embryos at dome stage. Cells were ordered from epiblast to ectoderm or mesoderm and endoderm, and colored by either pseudotime (left) or clusters (right).

**(D)** Bar chart showing numbers of upregulated gene (red), and downregulated gene (blue) at dome and shield stages in each cell cluster.

**(E)** Bubble plot showing enriched GO terms of differentially expressed genes (DEGs) between control and mKD embryos of each cluster at dome stage. Development and *p53* dependent apoptosis, immune response and cell cycle related terms are shown. Size of circle encodes gene number; color of circle indicates enrichment significance (-log10(P-value), modified Fisher's exact test).

Supplementary Figure 6

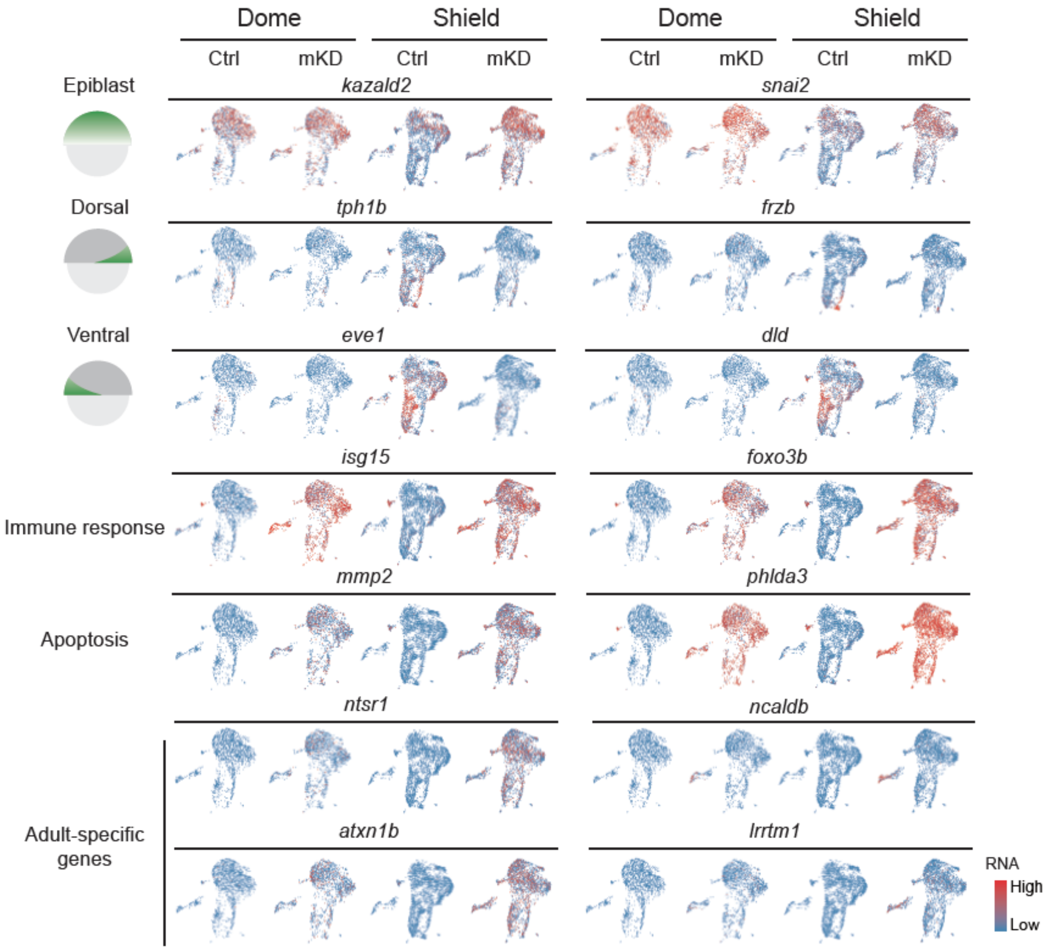


**Fig. S6. Aberrant transcription in *dnmt1* mKD embryos revealed by scRNA-seq.**
Feature plots showing RNA expression of marker genes of various lineages (models on the left) in control and *dnmt1* mKD embryos at dome and shield stages. Each dot represents one cell. Red means high expression; blue means low expression.

Supplementary Figure 7

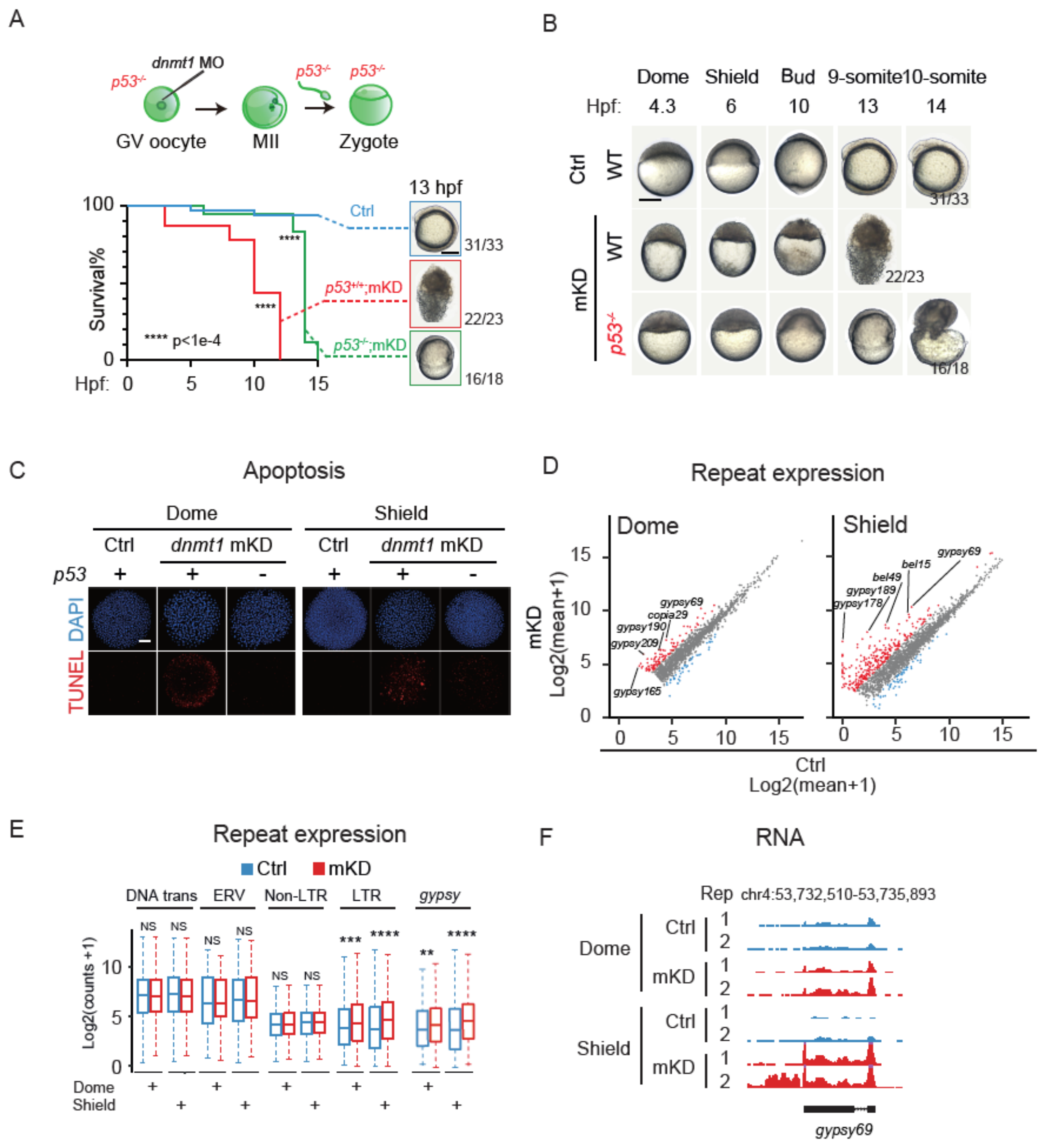


**Fig. S7.** ***dnmt1* mKD embryo lethality is in part caused by transposon-derepression triggered *p53*-mediated cell apoptosis.**

**(A)** Schematic of *dnmt1* mKD in $p53^{-/-}$ embryos (top). Survival curves of control (blue), *dnmt1* mKD in $p53^{+/+}$ (red) and *dnmt1* mKD in $p53^{-/-}$ (green) embryos are also shown. Representative images of embryo phenotype in each group are shown on the right. The numbers and ratios of embryos with this phenotype are shown. Scale bar, 250 µm.

**(B)** Representative images of embryo phenotype in each group at different stages. The numbers and ratios of embryos with corresponding phenotypes are also shown. Scale bar, 250 µm.

**(C)** TUNEL assay (red) of control, *dnmt1* mKD in $p53^{+/+}$ and $p53^{-/-}$ embryos at dome and shield stages. DNA was stained with DAPI (blue). Scale bar, 50 µm.

**(D)** Scatter plots comparing repeat expression (log2 (FPKM + 1)) between control and *dnmt1* mKD embryos at dome and shield stages. Red dots indicate upregulated repeats; blue dots indicate downregulated repeats.

**(E)** Box plots comparing RNA levels for four types of repeats (DNA trans, ERV, Non-LTR, and LTR) and *gypsy* repeat family at dome and shield stages of control (blue) and *dnmt1* mKD embryos (red). DNA trans, DNA transposons. ERV, endogenous retroviruses. Non-ERV, non-endogenous retroviruses. LTR, long terminal repeats. **, p<0.05, ****, p<0.0001, NS, not significant.

**(F)** A UCSC genome browser snapshot showing *gypsy69* expression at dome and shield stages of control (blue) and *dnmt1* mKD embryos (red).

Supplementary Figure 8

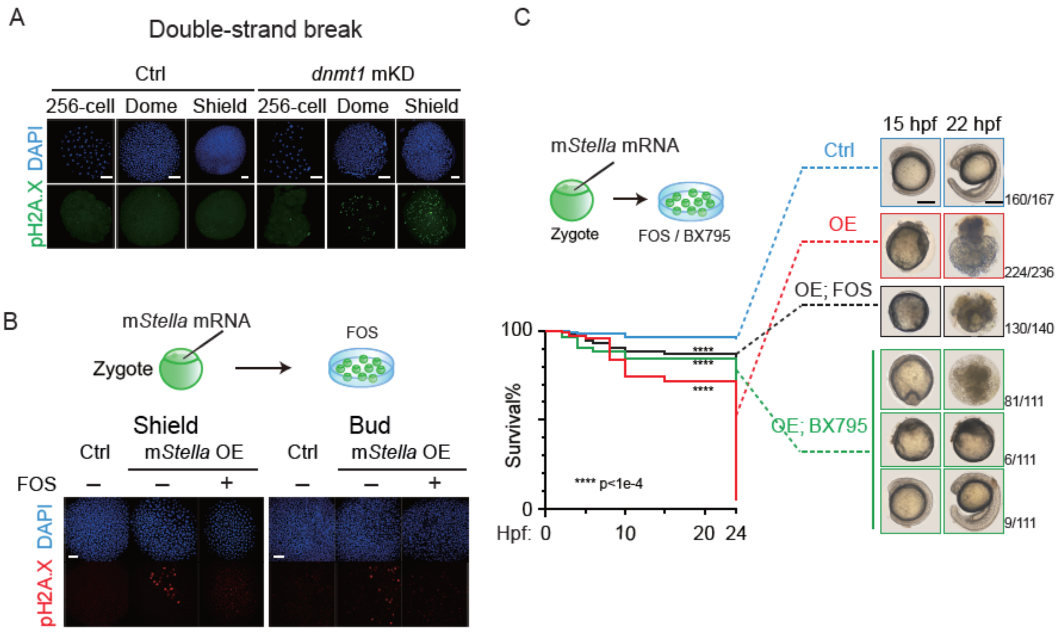


**Fig. S8. *dnmt1* mKD embryo lethality is in part caused by transposon-derepression triggered immune responses.**

**(A)** Immunostaining of pH2A.X (green) with control and *dnmt1* mKD embryos at 256-cell, dome and shield stages. DNA was stained with DAPI (blue). Scale bar, 50 µm.

**(B)** Schematic of *Stella* OE embryos treated with FOS. pH2A.X (red) immunostaining is shown for control, *Stella* OE and FOS treated *Stella* OE embryos at shield and bud stages. DNA was stained with DAPI (blue). Scale bar, 50 µm.

**(C)** Schematic of *Stella* OE embryos treated with FOS or BX795. Survival curves of control (blue), *Stella* OE (red), FOS treated *Stella* OE (black) and BX795 treated *Stella* OE (green) embryos are shown. Representative images of embryo phenotype in each group are shown on the right. The numbers and ratios of embryos with this phenotype are also shown. Scale bar, 250 µm.

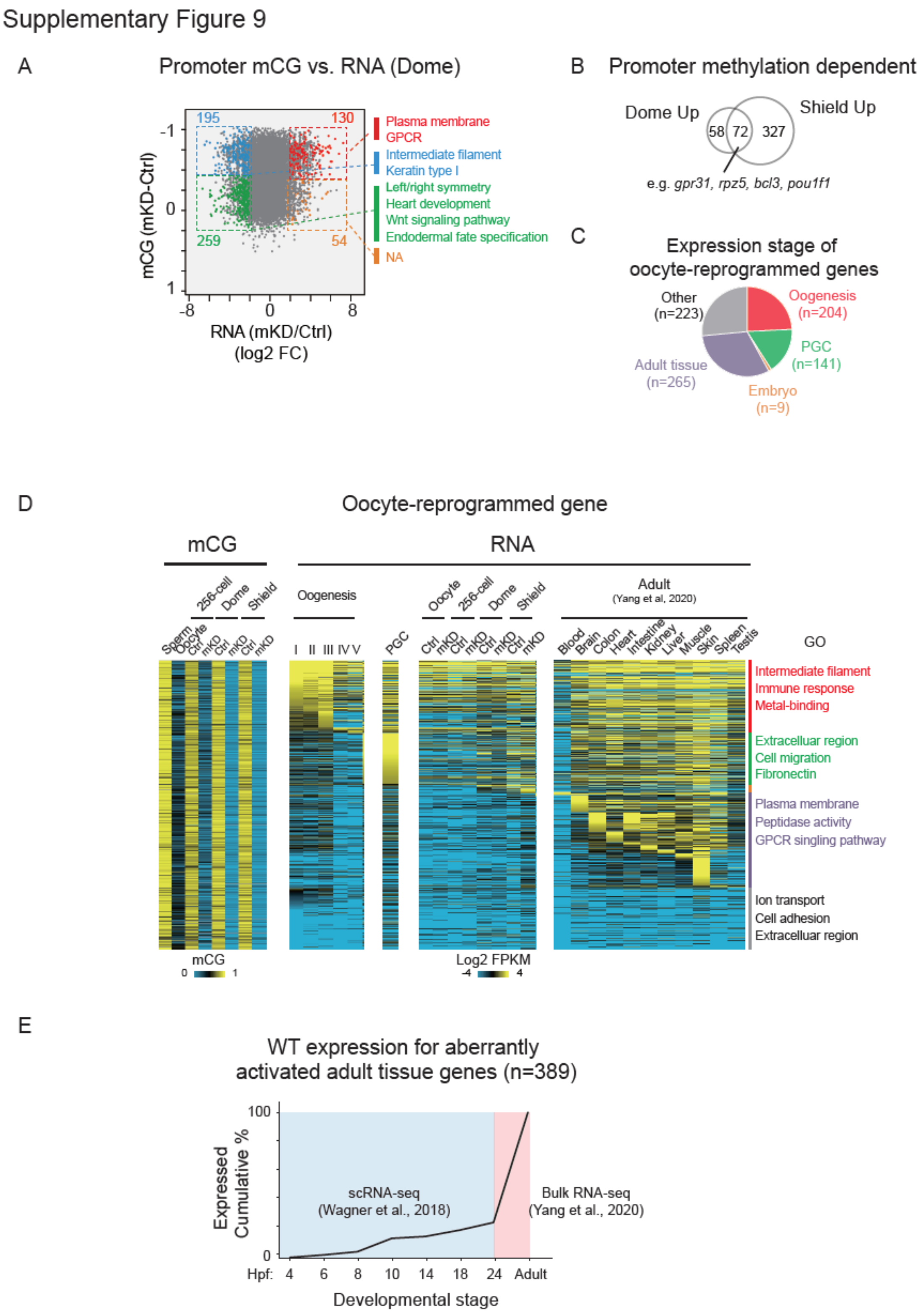


**Fig. S9. Relationship between DNA methylation and transcription in *dnmt1* mKD embryo.**
**(A)** Scatter plots comparing changes of gene expression (log2FC (mKD/Ctrl)) and promoter mCG between control and *dnmt1* mKD embryos at dome stage. Red and orange dots indicate promoter DNA methylation dependent and independent upregulated genes, respectively; blue and green dots

indicate downregulated genes with decreased and constant promoter DNA methylation, respectively. The numbers of dysregulated genes and enriched GO terms in corresponding group are also shown.
**(B)** Venn diagram showing overlap of upregulated genes upon the loss of DNA methylation between dome and shield stages.
**(C)** Pie chart showing distribution of expressing stages for all oocyte-reprogrammed genes (mCG (dome or shield - oocyte) > 0.4). The gene numbers in each cluster are shown.
**(D)** Heat maps showing mCG (left) and RNA (right) levels for all oocyte-reprogrammed genes. in sperm, oocytes during oogenesis, in 24hpf PGCs, control and *dnmt1* mKD oocytes, 256-cell, dome, shield stages and 24hpf embryos, larva, and in adult tissues (Yang et al., 2020). Genes derepressed in *dnmt1* mKD embryo were excluded. Enriched GO terms of these genes are shown on the right.
**(E**) Line chart showing among aberrantly activated adult expressed genes in *dnmt1* mKD embryos, what percentages (cumulated percentages) of genes are expressed by a particular stage in WT embryos. Data from 4hpf to 24hpf are based on published scRNA-seq data (Wagner et al., 2018). Data in adult tissue cells are described previously (Yang et al., 2020).

Supplementary Figure 10

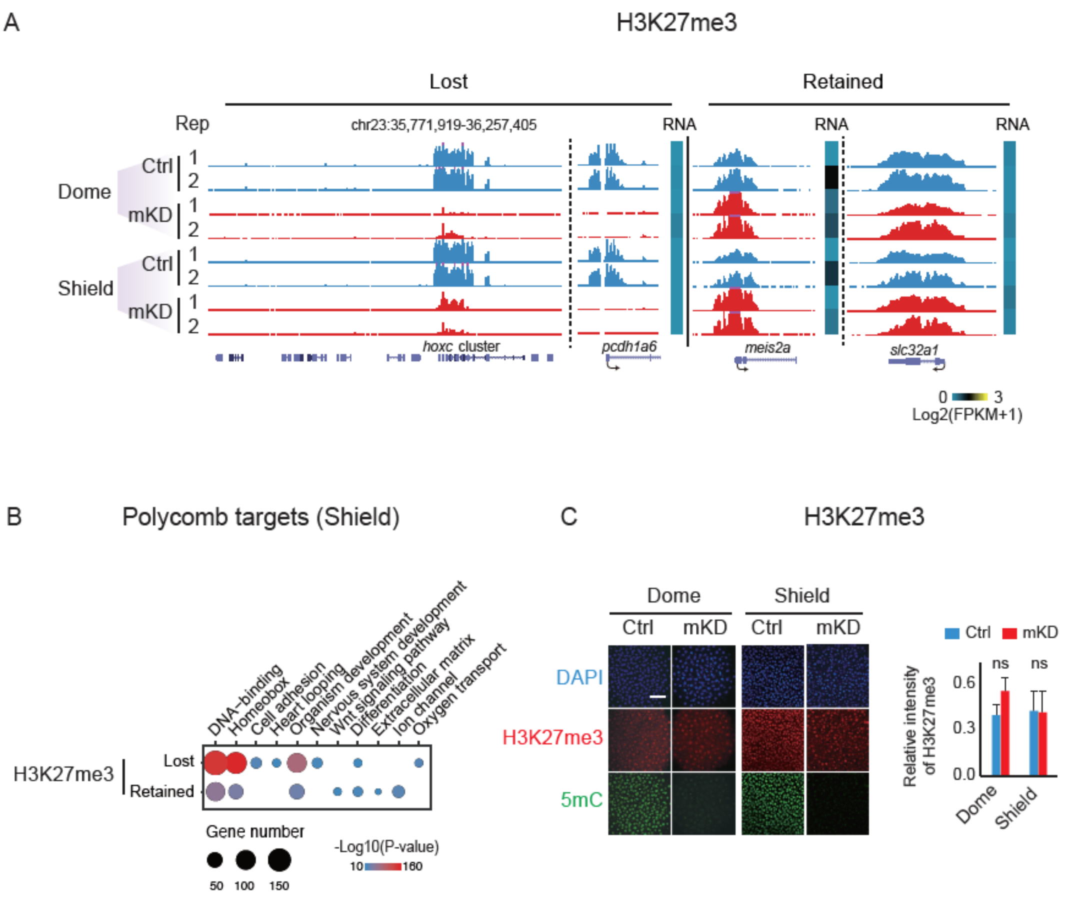


**Fig. S10. *dnmt1* mKD results in deficiency in H3K27me3 deposition in early embryos.**

**(A)** UCSC genome browser snapshots showing promoter H3K27me3 at dome and shield stages of control (blue) and *dnmt1* mKD embryos (red). Heat maps showing expression levels of these genes.

**(B)** Bubble plot showing enriched GO pathways of lost and retained promoter H3K27me3 at shield stage. Size of circle encodes gene number; color of circle indicates -log10 (P-value).

**(C)** Immunostaining of H3K27me3 (red) and 5mC (green) in control and *dnmt1* mKD embryos at dome and shield stages. The statistical evaluation was performed using student's t-test (control (blue) and *dnmt1* mKD (red)). DNA was stained with DAPI (blue). Scale bar, 50 µm.

### Supplementary Figure 11

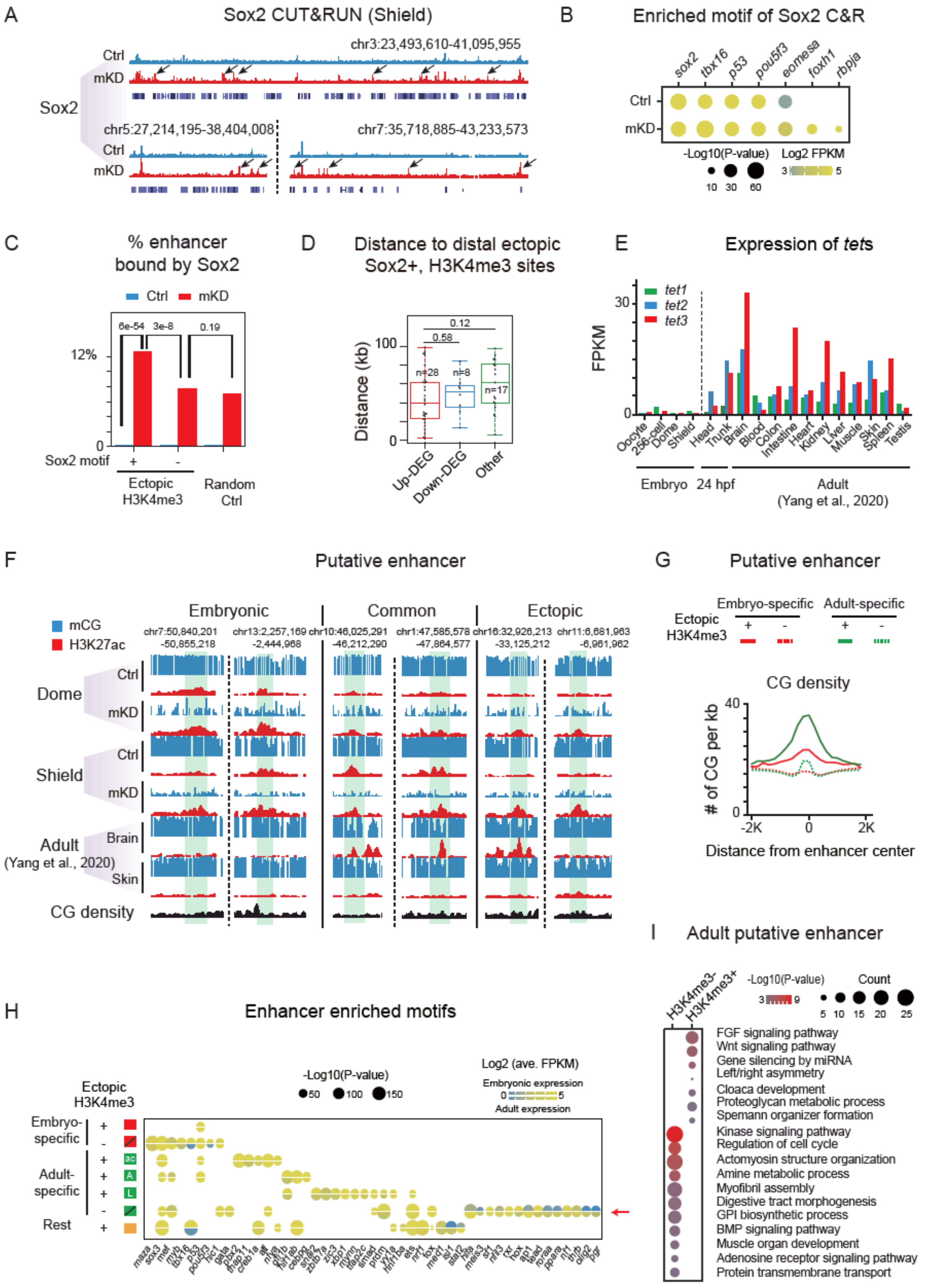


**Fig. S11. Adult enhancers are repressed by DNA methylation in early embryos.**

**(A)** UCSC genome browser snapshots showing Sox2 binding in control and *dnmt1* mKD embryos at shield stage. Black arrows indicate ectopic Sox2 binding.

**(B)** Bubble plot showing enriched motif of Sox2 binding sites in control and *dnmt1* mKD embryos at shield stage. Size of circle encodes -log10 (P-value), and color of circle indicates expression level of these genes at shield stage (log2FPKM).

**(C)** Bar chart showing fraction of enhancer bound by Sox2 at shield stage in control (blue) and *dnmt1* mKD (red) embryos. Somatic enhancers are randomly picked as random control.

**(D)** Box plot showing distance from center of Sox2 marked distal ectopic H3K4me3 peaks to upregulated (red), downregulated (blue), and non-DEGs (green) in *dnmt1* mKD embryos at shield stage.

**(E)** Bar chart showing expression of *tet* genes in embryos and adult tissues (Yang et al., 2020).

**(F)** UCSC genome browser snapshots showing mCG (blue), H3K27ac (red) and CG density (black) at enhancer regions in control and *dnmt1* mKD embryos at dome and shield stages, and in adult tissues brain and skin (Yang et al., 2020). Green shades indicate putative enhancers.

**(G)** Line plot showing CG densities around H3K4me3+ (light red) or H3K4me3- (dark red) embryonic enhancers, as well as H3K4me3+ (light green) or H3K4me3- (dark green) adult enhancer regions.

**(H)** Bubble plots showing TF motif enrichment in seven classes of putative enhancers. Color indicates expression (log2 (average FPKM)) levels in embryos (top) and adult tissues (bottom). Size indicates enrichment significance (-log10 (P-value)). The red arrow indicates a set of TFs that are preferentially expressed in adults but less so in embryos.

**(I)** Bubble plot showing GREAT analysis of H3K4me3+ and H3K4me3- adult enhancers. Size of circle encodes peak number; color of circle indicates -log10 (P-value).
